# Scalable, continuous evolution for the generation of diverse enzyme variants encompassing promiscuous activities

**DOI:** 10.1101/2020.06.01.128165

**Authors:** Gordon Rix, Ella J. Watkins-Dulaney, Patrick J. Almhjell, Christina E. Boville, Frances H. Arnold, Chang C. Liu

## Abstract

Enzyme orthologs sharing identical primary functions can have different promiscuous activities. While it is possible to mine this natural diversity to obtain useful biocatalysts, generating comparably rich ortholog diversity is difficult, as it is the product of deep evolutionary processes occurring in a multitude of separate species and populations. Here, we take a first step in recapitulating the depth and scale of natural ortholog evolution on laboratory timescales. Using a continuous directed evolution platform called OrthoRep, we rapidly evolved the *Thermotoga maritima* tryptophan synthase β-subunit (*Tm*TrpB) through multi-mutation pathways in many independent replicates, selecting only on*Tm*TrpB’s primary activity (synthesizing L-tryptophan from indole and L-serine). We find that the resulting sequence-diverse*Tm*TrpB variants span a range of substrate profiles useful in industrial biocatalysis and suggest that the depth and scale of evolution that OrthoRep affords will be generally valuable in enzyme engineering and the evolution of new biomolecular functions.

## Introduction

Natural enzymes typically have many orthologs. While the primary activity of orthologous enzymes is largely the same,^1^ promiscuous functions not under selective pressure can vary widely.^2,3^ Such variation may be attributed to the deep and distinct evolutionary histories shaping each ortholog, including long periods of neutral drift, recalibration of primary activity, and adaptation to new host environments such as temperature. These rich histories act to produce extensive genetic diversity, which underpins different promiscuity profiles.^2^

Diversity in promiscuous functions across orthologs is of both fundamental and practical importance. An enzyme’s reserve of promiscuous activities dictates what secondary reactions, environmental changes, or niches the enzyme can accommodate.^4,5^ Diversity in promiscuous activities therefore contributes to the basic robustness of life and adaptation. An enzyme’s reserve of promiscuous activities can also be mined in the application of enzymes for biocatalysis.^6,7^ Ortholog diversity therefore expands the range of reactions at the disposal of enzyme engineers, supporting the growing role of “green” enzymatic processes in the chemical and pharmaceutical industries.^8–10^

Inspired by the remarkable ability of enzyme orthologs to encompass promiscuous activities, we asked whether we could extend the substrate scope of useful enzymes by evolving multiple versions of an enzyme in the laboratory, selecting only for its primary function. Although this idea has been explored before using classical directed evolution approaches, most notably through the generation of cryptic genetic variation with neutral drift libraries,^11–14^ we recognized that our recently developed continuous evolution system, OrthoRep, may be considerably better poised for this challenge.^15,16^ Classical directed evolution mimics evolution through an iterative procedure that involves diversifying a gene of interest (GOI) *in vitro* (*e.g.*, through error-prone PCR), transforming the resulting GOI library into cells, and selecting or screening for desired activities, where each cycle of this procedure represents one step in an evolutionary search.^17^ However, since each cycle is manually staged, classical directed evolution does not readily admit depth and scale during exploration of functional sequence space — it is difficult to carry out many iterations to mimic lengthy evolutionary searches (depth), let alone do so in many independent experiments (scale). Yet evolutionary depth and scale are precisely the two characteristics responsible for ortholog diversity in nature. Natural orthologs have diversified from their ancestral parent over great evolutionary timescales, allowing for the traversal of long mutational pathways shaped by complex selection histories (depth). Natural orthologs are also the result of numerous independent evolutionary lineages, since spatially separated species and populations are free to take divergent mutational paths and experience different environments (scale). Systems that better mimic the depth and scale of natural enzyme evolution, but on laboratory timescales, are thus needed for the effective generation of enzyme variants that begin to approach the genetic and promiscuity profile diversity of orthologs.

OrthoRep is such a system. In OrthoRep, an orthogonal error-prone DNA polymerase durably hypermutates an orthogonal plasmid (p1) without raising the mutation rate of the host *Saccharomyces cerevisiae* genome.^16^ Thus, GOIs encoded on p1 rapidly evolve when cells are simply passaged under selection. By reducing the manual stages of classical directed evolution down to a continuous process where cycles of diversification and selection occur autonomously *in vivo*, OrthoRep readily accesses depth and scale in evolutionary search.^16,18^ Here, we apply OrthoRep to the evolution of the *Thermotoga maritima* tryptophan synthase β-subunit (*Tm*TrpB) in multiple independent continuous evolution experiments, each carried out for at least 100 generations. While we only pressured *Tm*TrpB to improve its primary activity of coupling indole and serine to produce tryptophan, the large number of independent evolution experiments we ran (scale) and the high degree of adaptation in each experiment (depth) resulted in a panel of variants encompassing expanded promiscuous activity with indole analogs. In addition to the immediate value of these newly evolved *Tm*TrpBs in the synthesis of tryptophan analogs, our study offers a new template for enzyme engineering where evolutionary depth and scale is leveraged on laboratory timescales to generate effective variant collections covering broad substrate scope.

## Results

### Establishing a selection system for the evolution of *Tm*TrpB variants

To evolve *Tm*TrpB variants using OrthoRep, we first needed to develop a selection where yeast would rely on *Tm*TrpB’s primary enzymatic activity for growth. *Tm*TrpB catalyzes the PLP-dependent coupling of L-serine and indole to generate L-tryptophan (Trp) in the presence of the tryptophan synthase α-subunit, *Tm*TrpA.^19^ In *T. maritima* and all other organisms that contain a heterodimeric tryptophan synthase complex, TrpA produces the indole substrate that TrpB uses and the absence of TrpA significantly attenuates the activity of TrpB through loss of allosteric activation.^19,20^ TRP5 is the *S. cerevisiae* homolog of this heterodimeric enzyme complex, carrying out both TrpA and TrpB reactions and producing Trp for the cell. We reasoned that by deleting the *TRP5* gene and forcing *S. cerevisiae* to rely on *Tm*TrpB instead, cells would be pressured to evolve high stand-alone *Tm*TrpB activity in order to produce the essential amino acid Trp in indole-supplemented media (Fig. 1a). This selection pressure would also include thermoadaptation, as yeast grow at mesophilic temperatures in contrast to the thermophilic source of *Tm*TrpB. Therefore, the selection on *Tm*TrpB’s primary activity would be multidimensional — stand-alone function, temperature, and neutral drift implemented when desired — and could result in complex evolutionary pathways that serve our goal of maximizing functional variant diversity across replicate evolution experiments. In addition, the multidimensional selection also serves practical goals as stand-alone activity is useful in biosynthetic applications (enzyme complexes are difficult to express and use *in vitro*) and activity at mesophilic temperatures is more compatible with heat-labile substrates, industrial processes where heating costs can compound, or *in vivo* applications in model mesophilic hosts (*e.g. S. cerevisiae* or *Escherichia coli*).

**Fig. 1.**
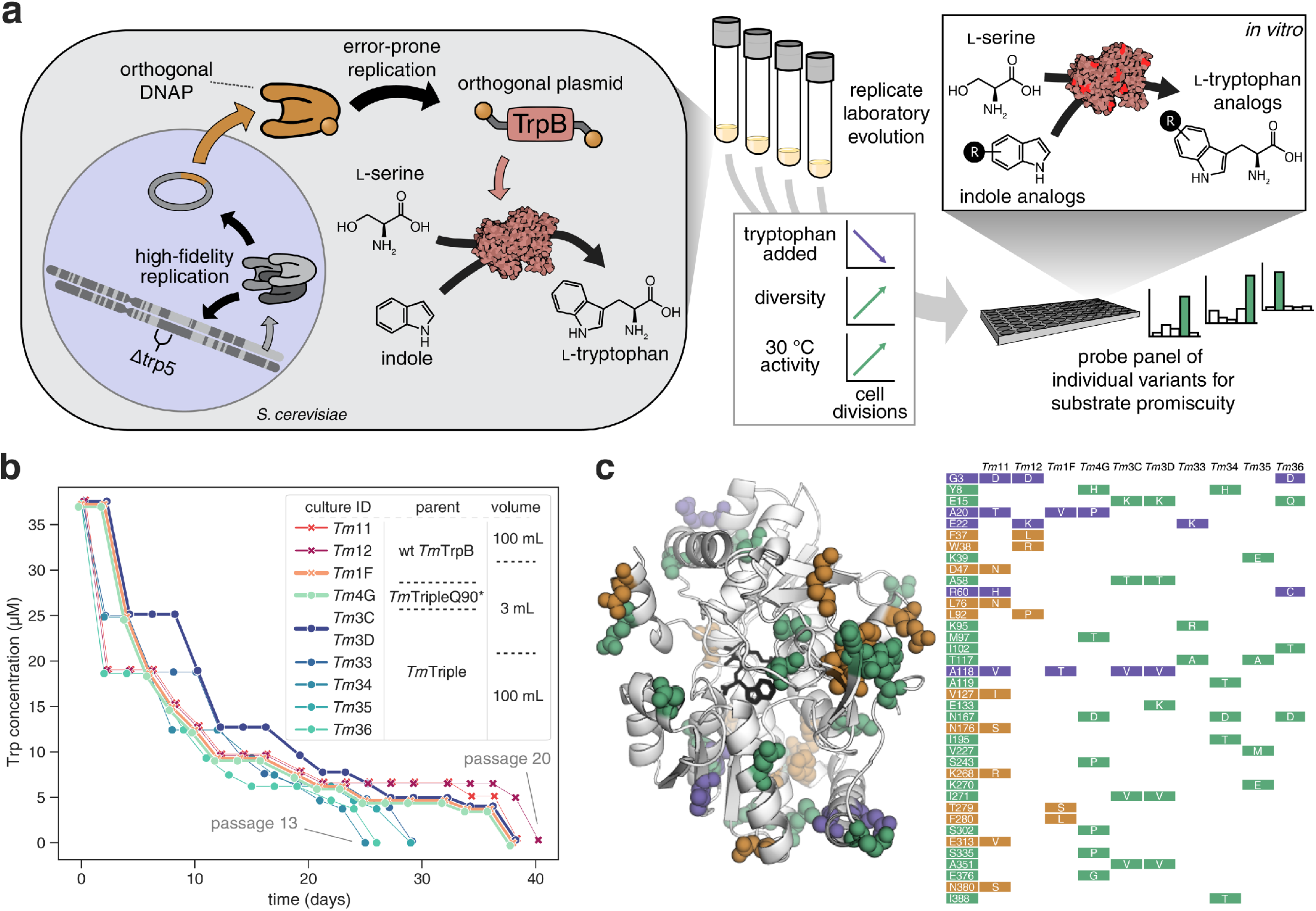
OrthoRep-mediated continuous *in vivo* evolution of *Tm*TrpB to generate many diverse, functional variants. **a**, Pipeline for the use of OrthoRep continuous directed evolution to generate many diverse, functional *Tm*TrpB sequences. *Tm*TrpB variants are first evolved in replicate for Trp production in yeast. OrthoRep enables replicate evolution through error-prone replication of an orthogonal plasmid by an orthogonal polymerase, maintaining low error rates in genome replication. By encoding a *Tm*TrpB variant on this plasmid in a tryptophan synthase (TRP5) deletion mutant, *Tm*TrpB may be both continuously diversified and selected for through gradual reduction in Trp supplied in the growth medium. Evolved populations containing many diverse, functional individuals may then be randomly sampled and tested for activity with indole analogs. *Tm*TrpB illustration generated using Illustrate.^35^ **b**, Selection trajectories for ten replicate cultures that evolved sufficient *Tm*TrpB activity to support cell growth without supplemented Trp. Each point represents a single 1:100 dilution (passage) into fresh indole-supplemented growth medium. Trp concentration of fresh media was reduced when high saturation was achieved in the previous passage. Plots are slightly offset from true values to allow for visibility of all selection trajectories. *Tm*3C and *Tm*3D are plotted as one line as their trajectories were identical. **c**, *Tm*TrpB homology model and table depicting consensus mutations of the ten cultures shown in panel **b**. Mutations are colored by their appearance in populations evolved from wt *Tm*TrpB (yellow), *Tm*Triple or *Tm*TripleQ90* (green), or both (purple).

To test this selection, we turned to a positive control *Tm*TrpB variant called *Tm*Triple. This variant was previously engineered to enable stand-alone activity, free from dependence on allosteric activation by TrpA, through a minimal set of three mutations.^7^ We found that *Tm*Triple rescued TRP5 function in a Δ*trp5* strain in an indole-dependent manner, validating our selection (Supplementary Fig. 1). Notably, *Tm*Triple, along with other TrpB variants tested, only supported complementation when expressed from a high-strength promoter (Supplementary Figs. 1a and 1b). This highlighted the opportunity for substantial adaptation and drift even in evolution experiments that start from already engineered *Tm*TrpB variants.

### Continuous evolution of *Tm*TrpB with depth and scale

We encoded wild-type (wt) *Tm*TrpB, *Tm*Triple, as well as a nonsense mutant of *Tm*Triple, *Tm*TripleQ90*, onto OrthoRep’s p1 plasmid, which is replicated by a highly error-prone orthogonal DNA polymerase. *Tm*TripleQ90* was included because reversion of the stop codon at position 90 in *Tm*TripleQ90* would act as an early indication that adaptation was occurring, giving us confidence to continue passaging our evolution experiments for several weeks to maximize evolutionary search depth. In all three OrthoRep Δ*trp5* strains, the initial *Tm*TrpB sequences enabled only minimal indole-dependent complementation (Supplementary Fig. 1c). This was expected for wt *Tm*TrpB, which has low stand-alone enzymatic activity and *Tm*TripleQ90*, which has a premature stop codon; and was unsurprising for *Tm*Triple, since *Tm*Triple displayed indole-dependent complementation only when artificially overexpressed (Supplementary Fig. 1a).

To continuously evolve *Tm*TrpB, we passaged cells encoding wt *Tm*TrpB, *Tm*Triple, or *Tm*TripleQ90* on OrthoRep in the presence of 100 μM indole while reducing the amount of Trp in the medium over time. In total, six 100 mL and twenty 3 mL cultures were passaged, each representing a single independent evolutionary trajectory. Passages were carried out as 1:100 dilutions where Trp concentrations were decreased in the (N+1)^th^ passage if cells grew quickly in the N^th^ passage, until Trp was fully omitted. All six of the 100 mL cultures, and four of the twenty 3 mL cultures fully adapted and were capable of robust growth in indole-supplemented media lacking Trp after 90– 130 generations (13–20 passages) (Fig. 1b, Supplementary Table 1). Populations that did not achieve growth in the absence of Trp still adapted, but stopped improving at ~5 µM supplemented Trp, suggesting a suboptimal local fitness maximum, though it is unknown if they would have adapted further had the experiment continued. Cultures that did adapt fully were passaged for an additional ~40 generations without increasing selection stringency to allow for accumulation of further diversity through neutral drift.

For each of the 10 fully adapted populations, we PCR-amplified and bulk-sequenced the *Tm*TrpB alleles on the p1 plasmid. Mutations relative to the parent *Tm*TrpB variant detected at >50% frequency in each population were deemed consensus mutations for that population, with the exception of reversion of the stop codon in populations evolving *Tm*TripleQ90*. This stop codon reversion occurred at 100% frequency in the relevant populations and was not counted in any subsequent analyses due to its triviality. An average of 5.6 (± 2.3 s.d.) and a range of 3–11 consensus amino acid changes per population were observed (Fig. 1c, Supplementary Table 2). Some of these mutations occurred at residues previously identified as relevant in conformational dynamics (*e.g.*, N167D and S302P).^20–22^ Most mutations observed, however, have not been previously identified in laboratory engineering experiments, suggesting that even the consensus of these populations explored new regions of *Tm*TrpB’s fitness landscape, doing so with diversity across replicates (Fig. 1c) that might translate to diversity in promiscuous activities across evolved variants.

### Evolved *Tm*TrpB variants improve Trp production *in vivo* and contain cryptic genetic variation

To ensure that evolved *Tm*TrpB variants, and not potential host genomic mutations, were primarily responsible for each population’s adaptation, we cloned individual *Tm*TrpBs into a standard low copy yeast nuclear plasmid under a promoter that approximates expression from p1,^23,24^ transformed the variants into a fresh Δ*trp5* strain, and tested for their ability to support indole-dependent growth in the absence of Trp (Figs. 2 and Supplementary Fig. 2). Sixteen *Tm*TrpB mutants were tested, representing one or two individual variants from each of the ten fully adapted populations. We found that 12 of the 16 TrpB variants complemented growth to a similar degree as TRP5 when supplemented with 400 μM indole, demonstrating substantial improvement over their wt *Tm*TrpB and *Tm*Triple parents (Fig. 2a).

**Fig. 2.**
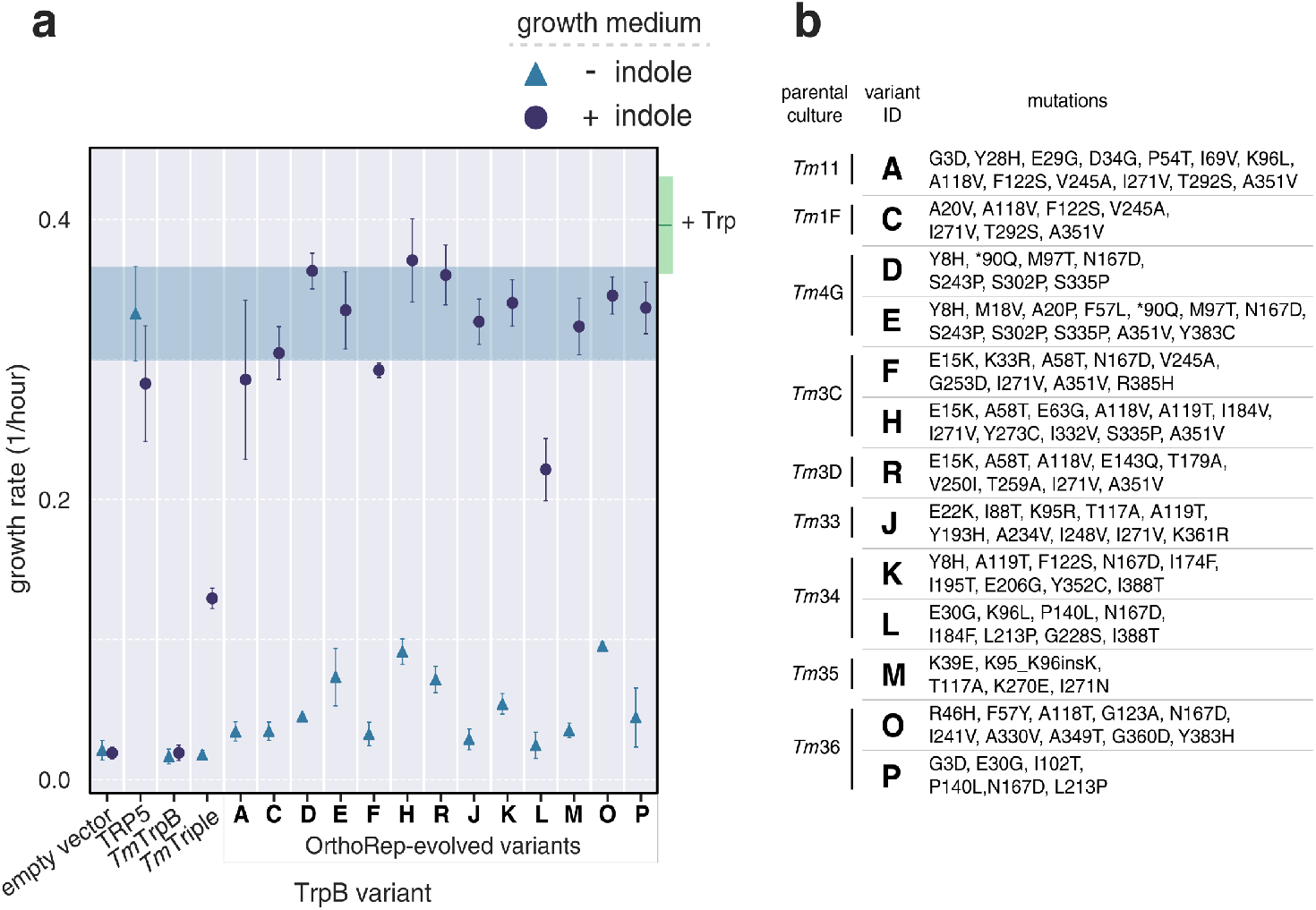
*In vivo* activity and diversity of individual *Tm*TrpB variants from OrthoRep-evolved populations. **a**, Evaluation of TRP5 complementation by evolved variants through a growth rate assay. Maximum growth rates over a 24-hour period for *Δtrp5* yeast strains transformed with a nuclear plasmid expressing the indicated *Tm*TrpB variant, grown in medium with or without 400 μM indole. Points and error bars represent mean ± s.d. for four biological replicates, respectively. Shaded area is the mean ± s.d. growth rate for the TRP5 positive control (*i.e.* plasmid expressing the endogenous yeast TRP5). Green box indicates the mean ± s.d. growth rate for all strains shown when Trp is supplemented. **b**, Parent populations from which OrthoRep-evolved variants shown in **a** are derived and all non-synonymous mutations present in each.

Unsurprisingly, this set of clonal *Tm*TrpBs contained more sequence diversity than the consensus sequences of the ten populations from which they were taken. Together, the variants tested comprised a total of 85 unique amino acid substitutions, with an average of 8.7 (± 2.1 s.d.) and a range of 5–13 non-synonymous mutations per variant (variant set 1, Supplementary Tables 2 and 3). Since the 12 *Tm*TrpBs from this set exhibiting complementation were all similarly active in their primary activity yet mutationally diverse (Fig. 2b), we may conclude that our scaled evolution experiments generated substantial cryptic genetic variation. We note that four of 16 *Tm*TrpB variants exhibited similar or lower Trp productivity compared to their parent (Supplementary Fig. 2b). We suspect that the multicopy nature of p1 in the OrthoRep system allowed for deleterious mutations to be maintained for a period of time without experiencing purifying selection if they arose in the same cell as functional variants, explaining the presence of these low activity *Tm*TrpBs. Indeed, this multicopy “buffering” may have worked to our advantage by promoting genetic drift under selection, facilitating both greater adaptation and greater diversity of evolutionary pathways across replicates (see **Discussion**). This may partly account for the high activity and high cryptic genetic variation present in the evolved *Tm*TrpBs.

### Evolved *Tm*TrpBs exhibit high primary and promiscuous activity *in vitro*

We further characterized the evolved *Tm*TrpBs *in vitro* to approximate conditions of industrial application, make kinetic measurements, and test whether promiscuous activity could be detected. Nine *Tm*TrpB variants were sampled from those that supported robust indole-dependent growth in the Δ*trp5* strain, cloned into an *E. coli* expression vector with a C-terminal polyhistidine tag, and overexpressed. To mimic streamlined purification conditions compatible with biocatalytic application of *Tm*TrpBs, we generated heat-treated *E. coli* lysates and tested them for their ability to couple indole and serine to produce Trp at 30 °C. Three of the nine OrthoRep-evolved TrpBs, **C**, **D**, and **K**, demonstrated improved activity over the benchmark *Tm*Triple, as measured by total turnover number (TTN) (Supplementary Fig. 3). Conveniently, each of these variants was evolved from a different starting point, meaning that wt *Tm*TrpB, *Tm*Triple, and *Tm*TripleQ90* were all viable starting points for reaching high activity *Tm*TrpBs.

Since the benchmark *Tm*Triple against which we compared the evolved *Tm*TrpBs was engineered through classical directed evolution involving screening *E. coli* lysates, whereas our *Tm*TrpB variants were evolved in yeast but expressed in *E. coli* for characterization, it is likely that the high-activity evolved *Tm*TrpBs would compare even more favorably if normalized by expression. We therefore purified **C**, **D**, and **K** by immobilized metal affinity chromatography (IMAC) and reevaluated their activity for coupling indole with serine to generate Trp. By TTN, all three variants showed a 4-to 5-fold increase in activity over *Tm*Triple at 30 °C (Supplementary Fig. 4a). At 75 °C, however, **C** had only ~2-fold higher activity than *Tm*Triple, while the other two variants were less active than *Tm*Triple. Since the thermostability of **C**, **D**, and **K** had not been reduced dramatically (T_50_ μ 83.7 °C, Supplementary Fig. 5), adaptation in **C**, **D**, and **K** occurred at least partially by shifting the activity temperature profile. This is a practically valuable adaptation, since thermostable enzymes that operate at mesophilic temperatures allow for greater versatility in application without sacrificing durability and ease of purification through heat treatment.

Further testing of **C**, **D**, and **K** revealed that all three enzymes had at least a 22-fold higher *k*_cat_/*K*_M_ for indole than did *Tm*Triple at 30 °C (Supplementary Table 4 and Supplementary Fig. 6). Finally, testing for production of Trp analogs revealed that these variants’ improved performance with indole transferred to alternate substrates (Supplementary Fig. 4b), validating their utility as versatile biocatalysts and also the hypothesis that continuous evolution of *Tm*TrpB variants can uncover promiscuous activities for which they were not selected.

### A diverse panel of evolved *Tm*TrpB variants encompasses a variety of useful promiscuous activities with indole analogs

Given the exceptional performance of **C**, **D**, and **K** and their ability to transfer primary activity to new substrates as promiscuous activity, we decided to further sample the variant diversity generated across the multiple *Tm*TrpB evolution experiments. We cloned 60 randomly chosen *Tm*TrpBs from the ten continuous evolution populations into *E. coli* expression vectors for *in vitro* characterization. These 60 *Tm*TrpBs represent extensive diversity, with an average of 9.3 (± 2.8 s.d.) non-synonymous mutations per variant and a total of 194 unique amino acid changes across the set; in addition, each sequence encoded a unique protein (variant set 2, Supplementary Tables 2 and 3). Since each variant had multiple non-synonymous mutations (up to 16) generated through >100 generations of adaptation and neutral drift, the depth of OrthoRep-based evolution was indeed leveraged in their evolution. We visualized these sequences, together with the consensus sequences of the populations from which they were derived, as nodes in a force directed graph related by shared mutations (Supplementary Fig. 7). With only one exception, all individual sequences cluster near the consensus sequence for their population, meaning that interpopulation diversity exceeded intrapopulation diversity. Thus, the scale of OrthoRep-based evolution was also leveraged in these variants — if fewer independent evolution experiments had been run, the reduction in diversity would not be recoverable from sampling more clones.

Preparations of *Tm*TrpBs **C**, **D**, and **K**, the 60 new variants, and four top-performing TrpB benchmark variants from past classical directed evolution campaigns (including *Tm*Triple) were all tested for product formation with indole by UV absorption and nine indole analogs by high performance liquid chromatography-mass spectrometry (HPLC-MS) to detect substrate promiscuity (Fig. 3a). The panel of 63 OrthoRep-evolved *Tm*TrpB variants exhibited an impressive range of activities (Fig. 3b). First, we observed that a number of variants had primary activities with indole that surpass the benchmark *Tm*Triple in lysate, with initial velocities of Trp formation up to 3-fold higher than **C**(Fig. 3b and Supplementary Fig. 8), whose *k*_cat_/*K*_M_ for indole is 1.37 × 10^5^ M^−1^ s^−1^, already 28-fold higher than *Tm*Triple’s at saturating serine concentrations (Supplementary Table 4 and Supplementary Fig. 6). Second, direct comparison of some of the best panel variants to *Tm*Triple revealed dramatic general activity improvements for multiple indole analogs (Fig. 3b). For example, across the three most versatile variants (**F7**, **D2**, and **F1**) the maximum fold-improvement in product yields over *Tm*Triple were 37, 5, 19, and 50 using substrates 5-cyanoindole (**2**), 7-cyanoindole (**4**), 5-bromoindole (**5**), and 6-bromoindole (**6**), respectively (Fig. 3c). Finally, with the exception of 6-bromoindole (**6**) and azulene (**10**), at least one variant from the OrthoRep-evolved panel converted the indole analog substrates as well as or better than benchmark TrpBs *Pf*2B9, *Tm*Azul, and *Tm*9D8*, which had been deliberately engineered toward new substrate scopes, though at higher temperatures (Figs. 3b and Supplementary Figs. 8 and 9).^7,21,25,26^

**Fig. 3.**
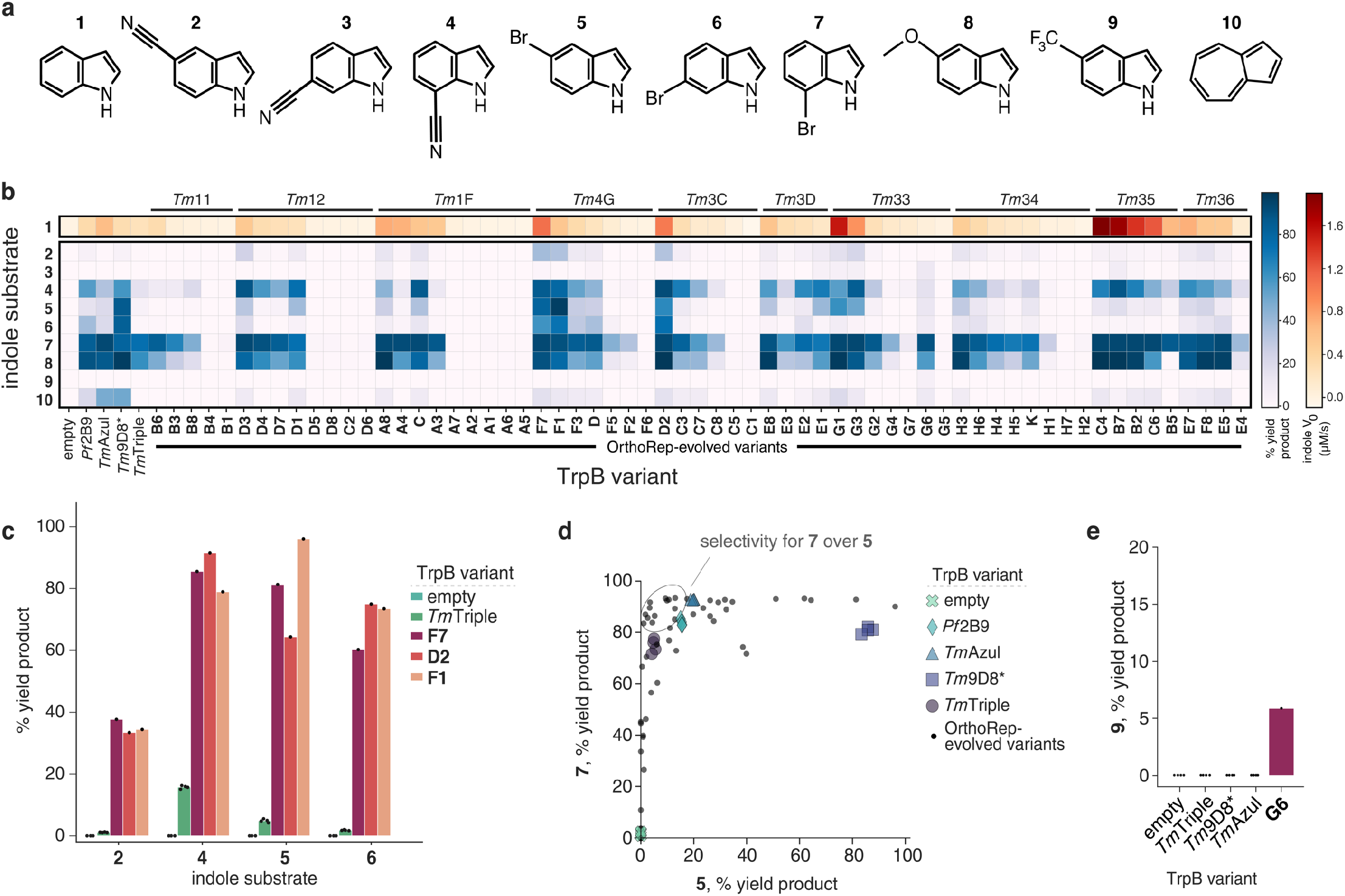
Promiscuous activities of a panel of evolved TmTrpBs. **a**, Indole substrates used to test the substrate scope of a panel of TmTrpB variants. **1**, indole; **2**, 5-cyanoindole; **3**, 6-cyanoindole; **4**, 7-cyanoindole; **5**, 5-bromoindole; **6**, 6-bromoindole; **7**, 7-bromoindole; **8**, 5-methoxyindole; **9**, 5-trifluoromethylindole; **10**, azulene. **b**, Heatmap of TrpB activities reported as yield of the Trp analog produced from indicated substrates where 100% yield corresponds to full conversion of the indole analog to the Trp analog. Reactions were carried out using heat-treated cell lysate, yield was measured by HPLC-MS, and V0 is the initial rate of Trp formation from indole at saturating serine concentrations. Panel TmTrpB variants are ordered first by the parental cultures from which they were derived (shown above the heatmap), then by activity with indole. Reactions with OrthoRep-evolved variants other than **C**, **D**, and **K** were performed in one replicate and all other reactions performed in quadruplicate. Empty designates expression vector without any TrpB encoded. **c**, Bar graph of selected indole analog activities from panel **b**. Points represent % HPLC yield for individual replicates, bars represent mean yield for multiple replicates or yield for a single replicate. **d**, Activities of all variants shown in **b** for reactions with substrates **7** and **5** to show selectivity. Individual replicates are shown for empty vector and benchmark TrpBs and only mean values are shown for OrthoRep-evolved variants tested in replicate (i.e. **C**, **D**, and **K**) for clarity. **e**, Heat treated lysate activity with **9** for indicated TrpB variants.

The diverse properties represented in our 63 variants were not just limited to primary activity increases on indole and promiscuous activities for indole analogs. Multiple variants from the panel also exhibited substantial improvements in selectivity for differently substituted indoles, which could be useful when working with substrate mixtures that may be less expensive to use industrially. For example, we observed many *Tm*TrpBs with greater selectivity for 7-bromoTrp over 5-bromoTrp as compared to all four of the benchmark engineered TrpBs (Fig. 3d). Another variant in the panel, **G6**, stood out for having appreciable activity with nearly all substrates tested, including 6-cyanoindole (**3**) and 5-trifluoromethylindole (**9**), which are poorly utilized by most other TrpBs tested, likely due to electron-withdrawing effects of their respective moieties. Notably, the ability to accept **9** as a substrate was unique to **G6**: all other variants, as well as the benchmark TrpBs, showed no detectable product formation with this substrate (Figs. 3e and Supplementary Fig. 9h). Repeating the reaction with purified enzyme validated the observed activity (Supplementary Fig. 10). **G6** may therefore be a promising starting point for new engineering efforts to access exotic Trp analogs. In short, despite having been selected for native activity with indole, OrthoRep-evolved *Tm*TrpBs have extensive and diverse activities on a range of non-native substrates, demonstrating the value of depth and scale in the evolution of enzyme variants.

### Mutations in evolved *Tm*TrpBs may modulate conformational dynamics and fine tune the active site

Of the ~200 unique mutations in the OrthoRep-evolved *Tm*TrpBs that we characterized, there were some mutations whose effects could be rationalized from comparison to previous work. Since these *Tm*TrpBs had to evolve stand-alone activity, it is unsurprising that many of the mutations we observed have been implicated in the loss of allosteric regulation by TrpA. For example, Buller *et al.* previously examined a series of engineered variants from *Pyrococcus furiosus* TrpB (*Pf*TrpB) and found that evolution for stand-alone activity was facilitated by a progressive shift in the rate-limiting step from the first to the second stage of the catalytic cycle as well as stabilization of the ‘closed’ conformation of the enzyme.^27^ That work implicated eight residues in this mechanism, seven of which correspond to homologous sites where we observed mutations in the evolved *Tm*TrpB variants (*i.e.*, P14, M18, I69, K96, L274, T292, and T321). Another mutation, N167D, present in three of the ten consensus sequences for evolved populations (Fig. 1c), has also been implicated in stabilizing the closed state.^21^ Additional mutations observed but not studied before (*e.g.*, S277F, S302P, and A321T) could also reasonably alter the allosteric network linking *Tm*TrpB activity to its natural *Tm*TrpA partner, based on existing structures and molecular dynamics simulations on the homologous *Pf*TrpA/*Pf*TrpB complex.^22,27^ Taken together, these mutations are likely implicated in converting allosteric activation by *Tm*TrpA into constitutive activity to establish stand-alone function of *Tm*TrpBs.

During the evolution of stand-alone activity, not only must allosteric activation by *Tm*TrpA be recapitulated by mutations in *Tm*TrpB, the surface of *Tm*TrpB that normally interacts with *Tm*TrpA must adjust to being exposed to solvent. Consistent with this adaptation, all consensus sequences for the ten successfully evolved populations from which our *Tm*TrpB variants were sampled contain a mutation to at least one of a set of five residues located on the canonical TrpA interaction interface (Fig. 1c and Supplementary Fig. 11).

We also detected strong convergent evolution in a region near the catalytic lysine, K83, which directly participates in *Tm*TrpB’s catalytic cycle through covalent binding of PLP and multiple proton transfers (Supplementary Fig. 12).^19^ For example, A118 was mutated in the consensus sequence of four of the ten fully adapted populations, while adjacent residues T117 or A119 were mutated in an additional three (Fig. 1c). Furthermore, the three populations in which these residues were not mutated contained other consensus mutations that are either part of the α-helix to which K83 belongs, or, like residues 117–119, within ~8 Å of this helix (Fig. 1c and Supplementary Fig. 12a). We hypothesize that the α-helix harboring K83 is a focal point of evolution, whereby mutations in its vicinity may finely adjust the positioning of K83 and the PLP cofactor to improve catalysis, perhaps as compensation for structural changes induced by thermoadaptation. Some OrthoRep-evolved variants also contained mutations to first- and second-shell active site residues (Supplementary Fig. 13), which may directly modulate the activity of *Tm*TrpBs, although these mutations were rare. Taken together, we hypothesize that these mutations near the active site residues of TrpB were adaptive or compensatory.

The ~20 mutations considered above are rationalized with respect to their impact on *Tm*TrpB’s primary catalytic activity. While substrate promiscuity changes may be influenced by these explainable mutations, previous literature suggests that substrate specificity is globally encoded by amino acids distributed across an entire enzyme.^28^ Indeed, the majority of the ~200 mutations found in our panel of *Tm*TrpBs were far away from *Tm*TrpB’s active site and not rationalizable based on the known structural and kinetic properties of TrpBs. We suspect that the cryptic genetic variation this majority of mutations encompasses contributes to the diversity in substrate scope across our variants.

## Discussion

In this work, we showed how the depth and scale of evolutionary search available in OrthoRep-driven protein evolution experiments could be applied to broaden the secondary promiscuous activities of *Tm*TrpB while only selecting on its primary activity. The significance of this finding can be divided into two categories, one concerning the practical utility of the new *Tm*TrpB variants we obtained and the second concerning how this evolution strategy may apply to future enzyme evolution campaigns and protein engineering in general.

Practically, the new *Tm*TrpBs should find immediate use in the synthesis of Trp analogs. Trp analogs are valuable chiral precursors to pharmaceuticals as well as versatile molecular probes, but their chemical synthesis is challenged by stereoselectivity requirements and functional group incompatibility. This has spurred enzyme engineers to evolve TrpB variants capable of producing Trp analogs,^20,21,25,26^ but the capabilities of available TrpBs are still limited. Compared to existing engineered TrpBs, our new panel of variants has substantially higher activity for the synthesis of Trp and Trp analogs at moderate temperatures from almost all indole analogs tested and also accepts indole analogs, such as 5-trifluoromethylindole, for which benchmark TrpBs used in this study showed no detectable activity (Fig. 3e). (In fact, only one TrpB variant has shown detectable activity for this substrate in previous classical directed evolution campaigns.^21^) In addition, at least one member of the panel accepted each of the nine indole analogs we used to profile promiscuity, suggesting that additional indole analogs and non-indole nucleophiles not assayed here will also be accepted as substrates.^29,30^ Finally, the evolved *Tm*TrpBs are both thermostable and adapted for enzymatic activity at 30 ΰC. This maximizes their industrial utility, as thermostability predicts a protein’s durability and can be exploited for simple heat-based purification processes, while mesophilic activity is compatible with heat-labile substrates, industrial processes where heating costs can compound, or *in vivo* applications in model mesophilic hosts (*e.g. S. cerevisiae* or *E. coli*).

Of more general significance may be the process through which the *Tm*TrpBs in this study were generated. Previous directed evolution campaigns aiming to expand the substrate scope of TrpB screened directly for activity on indole analogs to guide the evolution process,^21,26^ whereas this study only selected for *Tm*TrpB’s primary activity on indole. Yet this study still yielded *Tm*TrpBs whose secondary activities on indole analogs were both appreciable and diverse. Why?

A partial explanation may come from the high primary activities of OrthoRep-evolved *Tm*TrpBs, as validated by kinetic measurements showing that variants tested have *k*_cat_/*K*_M_ values for indole well in the 10^5^ M^−1^ s^−1^ range. Since OrthoRep drove the evolution of *Tm*TrpB in a continuous format for >100 generations, each resulting *Tm*TrpB is the outcome of many rounds of evolutionary improvement and change (evolutionary depth). This contrasts with previous directed evolution campaigns using only a small number of manual rounds of diversification and screening. Continuous OrthoRep evolution, on the other hand, allowed *Tm*TrpBs to become quite catalytically efficient with minimal researcher effort. We suggest that the high primary catalytic efficiencies also elevated secondary activities of *Tm*TrpB, resulting in the efficient use of indole analogs. However, this explanation is not complete, as evolved *Tm*TrpBs with similar primary activity on indole had differences in secondary activities (Fig. 3). In other words, high primary activities did not uniformly raise some intrinsic set of secondary activities in *Tm*TrpB, but rather influenced if not augmented the secondary activities of *Tm*TrpB in different ways. We attribute this to the fact that we ran our evolution experiments in multiple independent replicates (evolutionary scale). Each replicate could therefore evolve the same primary activity through different mutational paths, the idiosyncrasies of which manifest as distinct secondary activities. A third explanation for the promiscuous profile diversity of these *Tm*TrpB variants is that each replicate evolution experiment had, embedded within it, mechanisms to generate cryptic genetic variation without strong selection on primary activity. Many of the clones we sampled from each *Tm*TrpB evolution experiment had novel promiscuity profiles but mediocre primary activity with indole (Fig. 3). We believe this is because OrthoRep drove *Tm*TrpB evolution in the context of a multicopy plasmid such that non-neutral genetic drift from high activity sequences could occur within each cell at any given point. Therefore, *Tm*TrpB sequences with fitness-lowering mutations could persist for short periods of time, potentially allowing for the crossing of fitness valleys during evolution experiments and, at the end of each evolution experiment, a broadening of the genetic diversity of clones even without explicitly imposed periods of relaxed selection. Since enzyme orthologs are capable of specializing towards different sets of secondary activities when pressured to do so,2,3 non-neutral genetic drift from different consensus sequences across independent population should also access different secondary activities, further explaining the diversity of promiscuous activity profiles across clones selected from replicate evolution experiments. The combination of these mechanisms likely explains the variety of properties encompassed by the panel of *Tm*TrpBs.

Our approach to *Tm*TrpB evolution was inspired by the idea of gene orthologs in nature. Orthologs typically maintain their primary function while diversifying promiscuous activities through long evolutionary histories in different species.^2,31^ We approximated this by evolving *Tm*TrpB through continuous rounds of evolution, mimicking long histories, and in multiple replicates, mimicking the spatial separation and independence of species. Such depth and scale of evolutionary search is likely responsible for the substrate scope diversity of the *Tm*TrpBs we report even as they were selected only on their primary activity. We recognize that the evolved *Tm*TrpBs represent lower diversity than natural orthologs. For example, the median amino acid sequence divergence between orthologous human and mouse proteins is 11%,^32^ while the median divergence between pairs of variants from our experiment is 4.3% with a maximum of 8% (Supplementary Fig. 14). Still, this level of divergence between functional variants is substantial for a laboratory protein evolution experiment and suggests that it is realistic to model future work on the processes of natural ortholog evolution (Fig. 4). For example, it should be straightforward to scale our experiments further, to hundreds or thousands of independent populations each evolving over longer periods of time. This would better simulate the vastness of natural evolution. It should also be possible to deliberately vary selection schedules by adding competitive *Tm*TrpB inhibitors (such as the very indole analogs for which they have promiscuous activity), changing temperatures, or cycling through periods of weak and strong selection at different rates. Such evolutionary courses would approximate complexity in natural evolutionary histories. These modifications to OrthoRep-driven *Tm*TrpB evolution should yield greater cryptic genetic diversity, which may result in further broadening of promiscuous functions. The generation of cryptic genetic diversity at depth and scale should also be useful in efforts to predict protein folding and the functional effects of mutations via co-evolutionary analysis.^33,34^ Indeed, catalogs of natural orthologs have proven highly effective in fueling such computational efforts, so our ability to mimic natural ortholog generation on laboratory timescales may be applicable to protein biology at large. Within the scope of enzyme engineering, we envision that the process of continuous replicate evolution, selecting only on primary activities of enzymes, will become a general strategy for expanding promiscuous activity ranges of enzymes as we and others extend it to new targets.

**Fig. 4.**
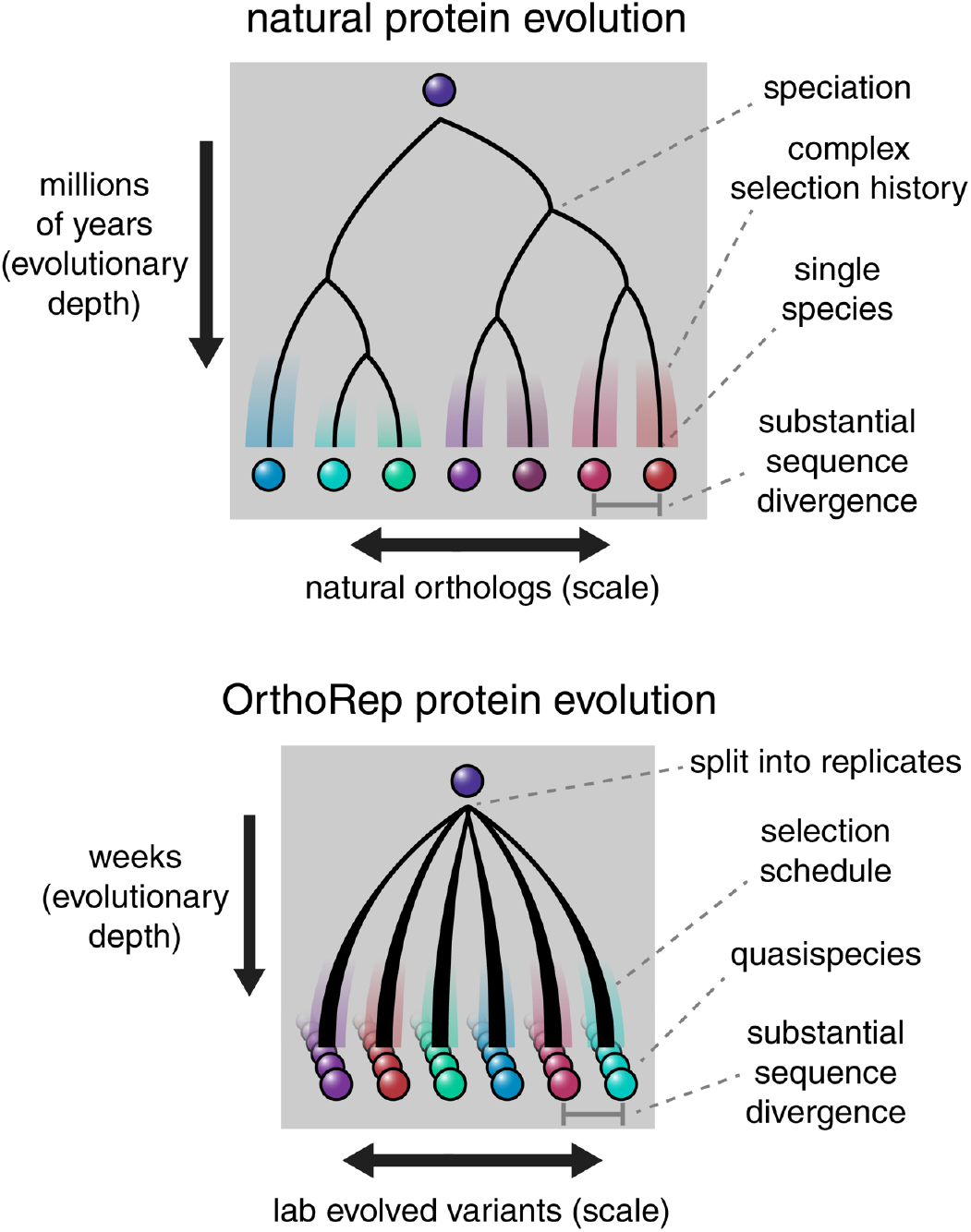
Conceptual similarities between natural enzyme ortholog evolution and OrthoRep evolution. Splitting OrthoRep cultures into many replicates can be seen as a form of speciation occurring by spatial separation. Complex selection schedules may emulate varied selection histories of natural orthologs, generating substantial sequence divergence across replicates. Evolved OrthoRep cultures contain diverse populations of sequences akin to quasispecies owing to high mutation rates.^36^

## Acknowledgments

This work was funded by NIH NIGMS (1DP2GM119163-01), NSF (MCB 1545158), and the Beckman Young Investigator Award to C.C.L., and the NIH NIGMS (R01GM125887) to F.H.A. The content is solely the responsibility of the authors and does not necessarily represent the official views of funding sources. We thank David Romney, Justin Bois, and other members of the Arnold and Liu groups for thoughtful discussions on experimental design.

## Author Contributions

All authors contributed to experimental design and data analysis. G.R. cloned all genetic constructs; set up, performed, and characterized evolution experiments; and carried out yeast growth rate experiments. E.J.W. performed the panel HPLC-MS assay and indole conversion rate measurements on *Tm*TrpBs from variant set 2, and P.J.A. analyzed the results. C.E.B. performed *in vitro* characterizations of *Tm*TrpBs from variant set 1 and performed the thermal shift assay and substrate scope characterizations for *Tm*TrpB variants **C**, **D**, and **K**. P.J.A. performed enzyme kinetics assays for *Tm*TrpB variants **C**, **D**, and G.R. and C.C.L. wrote the manuscript with input and contributions from all authors.

## Author information

*Christina E. Boville*

Present address: Aralez Bio, Emeryville, CA, USA

## Affiliations

*Department of Molecular Biology and Biochemistry, University of California, Irvine, Irvine, CA, USA* Gordon Rix, Chang C. Liu

*Department of Biomedical Engineering, University of California, Irvine, Irvine, CA, USA* Chang C. Liu

*Department of Chemistry, University of California, Irvine, Irvine, CA, USA* Chang C. Liu

*Division of Biology and Biological Engineering, California Institute of Technology, Pasadena, CA, USA* Ella J. Watkins-Dulaney, Frances H. Arnold

*Division of Chemistry and Chemical Engineering, California Institute of Technology, Pasadena, CA, USA* Patrick J. Almhjell, Frances H. Arnold

## Materials and methods

### DNA plasmid construction

Plasmids used in this study are listed in Supplementary Table 5. All plasmids that were not generated in a previous study were constructed via Gibson assembly^37^ from parts derived from the Yeast Toolkit,^24^ from previously described OrthoRep integration cassette plasmids,^16^ from *E. coli* expression vectors for previously described TrpB variants,^7,26^ from synthesized oligonucleotides, from yeast genomic DNA, or from the standard *E. coli* expression vector, pET-22b(+). All DNA cloning steps and *E. coli* protein expression steps were performed in *E. coli* strains TOP10 and BL21(DE3), respectively. All oligonucleotides used for PCR were purchased from IDT, and all enzymes and reagents used for cloning were purchased from NEB.

Parts used to generate yeast nuclear expression plasmids for testing the selection and p1 integration plasmids were PCR amplified from DNA sources listed above, Gibson assembled, transformed into *E. coli*, and plated onto selective LB agar plates. Individual clones were picked, grown to saturation in selective LB liquid media, miniprepped, and sequence confirmed. Following evolution of TrpB, individual variants were assembled into new yeast or *E. coli* expression vectors through PCR amplification of purified DNA from evolved yeast cultures, bulk cloning into the appropriate expression vector, picking individual colonies, and confirming absence of any frameshift mutations by Sanger sequencing.

### Yeast strains and media

All yeast strains used in this study are listed in Supplementary Table 6. Yeast were incubated at 30 °C, with shaking at 200 rpm for liquid cultures, and were typically grown in synthetic complete (SC) growth medium (20 g/L dextrose, 6.7 g/L yeast nitrogen base w/o amino acids (US Biological), 2 g/L SC dropout (US Biological) minus nutrients required for appropriate auxotrophy selection(s)), or were grown in YPD growth medium (10 g/L bacto yeast extract, 20 g/L bacto peptone, 20 g/L dextrose) with or without antibiotics, if no auxotrophic markers were being selected for. Media agar plates were made by combining 2X concentrate of molten agar and 2X concentrate of desired media formulation. Prior to all experiments, cells were grown to saturation in media selecting for maintenance of any plasmids present.

### Yeast transformation

All yeast transformations were performed as described in Gietz and Shiestl.^38^ After all transformations, transformed cells were streaked onto selective media agar plates, and resulting single colonies were picked for all further uses. Transformations for integration onto p1 were performed as described previously:^15^ 2–4 µg of plasmid DNA with ScaI restriction sites adjacent to integration flanks was cut with ScaI-HF (NEB) and transformed into yeast harboring the wt p1 and p2 plasmids. Proper integration was validated by miniprepping resulting clonal strain as previously described,^15^ visualizing the recombinant p1 band of the desired size by gel electrophoresis, and PCR and Sanger sequencing of the gene of interest integrated onto p1. Resulting strains were then transformed with either of two plasmids for nuclear expression of an OrthoRep terminal protein DNA polymerase 1 (TP-DNAP1) variant: wt TP-DNAP1 (pAR-Ec318) for evaluating *trp5* complementation of TrpB variants without mutagenesis, or error-prone TP-DNAP1 (pAR-Ec633) for generating strains ready for TrpB evolution. These strains were passaged for ~40 generations in order to stabilize copy number of the recombinant p1 species, prior to any use in experiments.

Genomic deletions were made through co-transformation of a CRISPR/Cas9 plasmid targeting the region of interest and a linear DNA fragment comprised of two concatenated 50 bp homology flanks to the region of interest.^39^ Transformations were then plated on selective media agar, colonies were re-streaked onto nonselective media agar, and resulting colonies were grown to saturation in liquid media. The region of interest was PCR amplified and Sanger sequenced to confirm presence of desired modification.

### Plating assays

Yeast strains expressing a TrpB variant either from a nuclear plasmid, or from p1, with wt OrthoRep polymerase (TP-DNAP1) expressed from a nuclear plasmid, were grown to saturation in SC –L or SC –LH, spun down, washed once with 0.9% NaCl, then spun down again, and the resulting pellet was resuspended in 0.9% NaCl. Washed cells were then diluted 1:100 in 0.9% NaCl, and 10 µL of each diluted cell suspension was plated onto media agar plates in pre-marked positions. After 3 days of growth, cell spots were imaged (Bio-Rad ChemiDoc™). Resulting images were adjusted uniformly (‘High’ set to 40,000) to improve visibility of growth (Bio-Rad Image Lab™ Software). Figures utilizing these images (Supplementary Figs. 1 and 2a) were made by manually combining images of different plates, but all images of the same media condition within each figure panel were derived from the same image of a single plate.

### *Tm*TrpB evolution

Yeast strains with a nuclear plasmid expressing error-prone TP-DNAP1 and with wt *Tm*TrpB, *Tm*Triple, or *Tm*TripleQ90* encoded on p1 (GR-Y053, GR-Y055, and GR-Y057, Supplementary Table 6) were grown to saturation in SC –LH, prior to passaging for evolution. All cultures passaged for evolution of *Tm*TrpB regardless of success are described in Supplementary Table 1. To provide enough indole substrate for sufficient Trp production, but not enough to induce toxicity, all growth media used for evolution of TrpB activity was supplemented with 100 μM indole, as informed by results shown in Supplementary Fig. 1a. All passages for evolution were carried out as 1:100 dilutions. In order to induce a growth defect but still allow for some growth, the first passage for each evolution culture was carried out in SC –LH media with 37 µM Trp (7.6 mg/L). After two or three days of shaking incubation, if OD_600_ μ 1.0 (Bio-Rad SmartSpec™ 3000) for 100 mL cultures, or if most wells in a 24 well block of 3 mL cultures were saturated to a similar degree by eye, cultures were passaged into fresh growth medium with a slightly reduced Trp concentration. If the level of growth was beneath this threshold, the culture was passaged into growth medium with the same Trp concentration. This process was continued until a Trp concentration of 3.7 µM was reached, at which point a passage into media lacking Trp was attempted, which typically resulted in successful growth. Resulting cultures were then passaged six additional times into growth medium lacking Trp.

### Growth rate assays

Yeast strains containing nuclear plasmids encoding one of several OrthoRep-evolved TrpB variants, wt *Tm*TrpB, *Tm*Triple, or none of these (denoted ‘empty’) were grown to saturation in SC –L, washed as described above, then inoculated 1:100 into multiple media conditions in 96-well clear bottom plates, with four biological replicates per media/strain combination. Plates were then sealed with a porous membrane and allowed to incubate with shaking at 30 °C for 24 hours, with OD_600_ measurements taken automatically every 30 minutes (Tecan Infinite M200 Pro), according to a previously described protocol.^40^ Multiple 24 hour periods were required for each experiment, but empty controls were included in each individual 96-well plate to ensure validity of growth in other cultures. Raw OD_600_ measurements were fed into a custom MATLAB script,^18^ which carries out a logarithmic transformation to linearize the exponential growth phase, identifies this growth phase, and uses this to calculate the doubling time (T). Doubling time was then converted to growth rate by the equation *ln(2)/T*.

### Enzyme characterization— general experimental methods

Chemicals and reagents were purchased from commercial sources and used without further purification. All cultures were grown in Terrific Broth supplemented with 100 Δg/mL carbenicillin (TBcarb). Cultures were shaken in a New Brunswick Innova 4000 (shaking diameter 19 mm), with the exception of the 96-well deep-well plates (USA Scientific), which were shaken in a Multitron INFORS HT (shaking diameter 50 mm). Lysis buffer was composed of 50 mM potassium phosphate, pH 8.0 (KPi buffer), supplemented with 100 or 200 µM pyridoxal 5’-phosphate (PLP). Heat lysis was performed in a 75 °C water bath (Fisher) for >1 hour. Protein concentrations were determined using a Pierce™ BCA Protein Assay Kit (Thermo Scientific). Reactions were performed in KPi buffer. Liquid chromatography/mass spectrometry (LCMS) was performed on an Agilent 1290 UPLC-LCMS equipped with a C-18 silica column (1.8 μM, 2.1 × 50 mm) using CH_3_CN/H_2_O (0.1% acetic acid by volume): 5% to 95% CH_3_CN over 2 min; 1 mL/min. Liquid chromatography/mass spectrometry (LCMS) was also performed on an Agilent 1260 HPLC-MS equipped with Agilent InfinityLab Poroshell 120 EC-C18 column (2.7 μM, 4.6×50 mm): hold 5% CH_3_CN for 0.5 min, 5-95% CH_3_CN over 2 min; 1 mL/min.

TrpB variants selected for further characterization were cloned into a pET-22b(+) vector with a C-terminal 6X His-tag and transformed into *E. coli* BL21(DE3) cells (Lucigen).

### Expression and characterization of variants from set 1 — large scale expression and lysis

A single colony containing the appropriate TrpB gene was used to inoculate 5 mL TB_carb_ and incubated overnight at 37 °C and 230 rpm. For expression, 0.5 mL of overnight culture were used to inoculate 50 mL TB_carb_ in a 250 mL flask and incubated at 37 °C and 250 rpm for 3 hours to reach OD_600_ 0.6–0.8. Cultures were chilled on ice for 20 min and expression was induced with a final concentration of 1 mM isopropyl β-D-thiogalactopyranoside (IPTG). Expression proceeded at 25 °C and 250 rpm for approximately 20 hours. Cells were harvested by centrifugation at 5,000*g* for 5 min at 4 °C, and then the supernatant was decanted. The pellet was stored at −20 °C until further use or used immediately for whole cell transformations.

Pellets were lysed in 5 mL of lysis KPi buffer with 200 μM PLP, supplemented with 1 mg/mL lysozyme (HEWL, Sigma Aldrich), 0.02 mg/mL bovine pancreas DNase I, and 0.1X BugBuster (Novagen) and incubated at 37 °C for 30 minutes. Lysate was clarified by centrifugation at 5,000*g* for 10 min, divided into 1 mL aliquots, and stored at −20 °C until further use.

### Expression and characterization of variants from set 1— lysate and whole cell small-scale reactions

Protein concentration in lysate was quantified by BCA. Lysate reactions were performed in 2 mL glass HPLC vials (Agilent) charged with indole (final conc. 20 mM) dissolved in DMSO (5% w/v), followed by the addition of lysate (final enzyme conc. 4 μM), and serine (final conc. 20 mM) to achieve a final volume of 200 μL. Whole cell reactions were performed in 2 mL glass HPLC vials (Agilent) charged with indole (final conc. 20 mM) dissolved in DMSO (5% w/v), followed by the addition of cells diluted in KPi buffer (final OD_600_=6), and serine (final conc. 20 mM) to achieve a final volume of 200 μL. Reactions were incubated at 30 °C for 24 hours, diluted with 800 μL 1:1 CH_3_CN/1 M aq. HCl, and analyzed via UHPLC-MS.

### Expression and characterization of variants from set 1— thermostability determination

Enzyme T_50_ measurements (the temperature at which 50% of the enzyme is irreversibly inactivated after a 1 hour incubation) were used to report on the thermostability of the enzyme. In a total volume of 100 µL, samples were prepared in KPi buffer with 1 µM enzyme in PCR tubes and either set aside (25 °C) or heated in a thermal cycler on a gradient from 79–99 °C (OrthoRep-generated variants), or 59–99 °C (*Tm*Triple), for 1 hour, with each temperature performed in duplicate. Precipitated protein was pelleted via centrifugation and 75 µL of each sample was carefully removed and added to the wells of a 96-well UV-transparent assay plate containing 0.5 mM indole and 0.5 mM serine. Relative product formation was observed by measuring the change in absorbance at 290 nm to determine the temperature at which the sample had 50% residual activity compared to the 25 °C samples (modeled as a logistic function).

### Expression and characterization of variants from set 1— enzyme kinetics

Enzymatic parameters, *k*_cat_ and *K*_M,_ for the conversion of indole to Trp were estimated via Bayesian inference (pystan 2.19.0.0) assuming Michaelis-Menten behavior under saturating serine (40 mM) in KPi buffer. Briefly, initial velocities (*v*) were determined by monitoring Trp formation in a Shimadzu UV-1800 spectrophotometer at 30 °C for 1 min over a range of indole concentrations at 290 nm using the reported indole-Trp difference in absorbance coefficient (Δε_290_ = 1.89 mM^−1^ cm^−1^).^41^ These velocities were modeled using the equation:

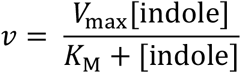

and estimates for *v* and *V*_max_ were converted to *k* and *k*_cat_ by normalizing for enzyme concentration. Parameter estimates are obtained as Hamiltonian Markov chain Monte Carlo (MCMC) posterior samples and reported as the median with their 95% credible regions (CR). The code used to generate these estimates (along with example data) can be found at http://github.com/palmhjell/bayesian_kinetics.

### Expression and characterization of variants from set 2 — small-scale expression and lysis

Variants were arrayed into a 96-well deep-well plate along with *Tm*Triple, *Tm*9D8*, *Tm*Azul, and *Pf*2B9. Individual colonies were grown in 600 μL TB_carb_ in 96-well polypropylene plates overnight at 37 °C, 250 rpm, 80% humidity. The following day, 20 μL of overnight culture was used to inoculate 630 μL TB_carb_ in deep-well 96-well plates and grown at 37 °C, 250 rpm. After 4 hours, cultures were chilled on ice for 20–30 min and protein expression was induced with 50 μL IPTG (final conc. 1 mM) diluted in TB_carb_. Cultures were shaken at 20 °C, 250 rpm for 20–24 hours, after which they were subjected to centrifugation at 5,000*g* for 10 min. The cell pellets were frozen at −20 °C until further use or used immediately.

### Expression and characterization of variants from set 2 — indole rate measurements

Pellets were lysed in either 600 μL of KPi buffer with 100 μM PLP and heat-treated at 75 °C for 1 hour, or in 600 μL of this buffer supplemented with 1 mg/mL lysozyme, 0.02 mg/mL bovine pancreas DNase I, and 0.1X BugBuster and incubated at 37 °C for 1 hour. Lysate from both conditions was clarified by centrifugation at 4,500*g* for 10 min and stored at 4 °C until further use.

Reaction master mix composed of 625 μM indole and 25 mM serine in KPi buffer was prepared and, before reactions, plates and master mix were incubated in 30 °C water bath for 30 min. The microplate reader (Tecan Spark) was also pre-heated to 30 °C.

To UV-transparent 96-well assay plates (Caplugs, catalog # 290-8120-0AF), 160 μL pre-heated reaction master mix was added by 12-channel pipet followed by 40 μL of lysate from the pre-heated plate using a Microlab NIMBUS96 liquid handler (Hamilton). Plates were immediately transferred into the plate reader, shaken for 10 sec to mix and the absorbance of each well at 290 nm was recorded as rapidly as possible (~20 sec between measurements) for 120 cycles. The rate of product formation was determined by finding the rate of absorbance change over time and converting to units of concentration using Δε_290_ = 1.89 mM^−1^ cm^−1^ (see above) and a determined path length of 0.56 cm. Differences between the two lysate preparations were not significant suggesting that most enzyme variants retained sufficient thermostability for purification via heat treatment, and this method was used in subsequent experiments.

### Expression and characterization of variants from set 2 — substrate scope screen

Pellets were lysed in 300 μL KPi buffer with 200 μM PLP and clarified by centrifugation at 4,000*g* for 10 min. To a 96-well deep-well plate charged with 10 μL nucleophile dissolved in DMSO (final conc. denoted in the table directly below), 40 μL of the heat-treated lysate was transferred using a Microlab NIMBUS96 liquid handler (Hamilton), followed by addition of 150 μL serine (final conc. 20 mM) with a 12-channel pipet. Reactions were sealed with 96-well ArctiSeal™ Silicone/PTFE Coating (Arctic White) and incubated in 30 °C water bath for ~24 hours. Reactions were diluted with 600 μL 2:1 CH_3_CN/1 M aq. HCl, subjected to centrifugation at 5,000*g*, and 400 μL was transferred to 2 mL glass HPLC vials (Agilent). Samples were analyzed by HPLC-MS. Azulene samples were further diluted 20X to avoid oversaturation of the UV-detector and analyzed via UHPLC-MS.

| Nucleophile | Source | Catalog # | CAS | Final conc. (mM) |
| --- | --- | --- | --- | --- |
| 5-Trifluoromethylindole | Sigma | 701068 | 100846-24-0 | 10 |
| 4-Cyanoindole | Sigma | 645532 | 100846-24-0 | 20 |
| 5-Cyanoindole | Chem Impex | 21849 | 15861-24-2 | 20 |
| 6-Cyanoindole | Chem Impex | 21181 | 15861-36-6 | 20 |
| 7-Cyanoindole | Sigma | CDS008484 |  | 20 |
| 5-Bromoindole | Sigma | B68607 | 10075-50-0 | 20 |
| 5-Methoxyindole | Combi-Blocks | IN0049 | 1006-94-6 | 20 |
| Azulene | Alfa Aesar | L08271 | 275-51-4 | 20 |

All samples except those containing azulene were analyzed at 277 nm, representing the isosbestic point between indole and Trp and allowing estimation of yield by comparing the substrate and product peak areas for indole analogs.^21^ Azulene yield was estimated as described previously.^25^ Nucleophile retention times were determined though injection of authentic standards and product retention times were identified by extracting their expected mass from the mass spectrum.

### Large-scale expression and purification of B5 and G6

A single colony containing the appropriate TrpB gene was used to inoculate 5 mL TB_carb_ and incubated overnight at 37 °C and 230 rpm. For expression, 2.5 mL of overnight culture were used to inoculate 250 mL TB_carb_ in a 1-L flask and incubated at 37 °C and 250 rpm for 3 hours to reach OD_600_ 0.6–0.8. Cultures were chilled on ice for 20 min and expression was induced with a final concentration of 1 mM IPTG. Expression proceeded at 25 °C and 250 rpm for approximately 20 hours. Cells were harvested by centrifugation at 5,000*g* for 5 min at 4 °C, and then the supernatant was decanted. The pellet was stored at −20 °C until further use.

Pellets were lysed in 25 mL KPi buffer with 200 μM PLP for >1 hour at 75 °C. Lysate was clarified by spinning 14,000*g* for 20 min at 4 °C (New Brunswick Avanti J-30I). Protein was purified over hand-packed HisPur™ Ni-NTA Resin (Thermo Scientific, catalog # 88221), dialyzed into KPi buffer and quantified by BCA.

### B5 PLP-binding assay

Variant B5 did not the exhibit characteristic yellow color of PLP-bound TrpB variants after purification, however BCA indicated comparable protein concentrations to the G6 variant. We have previously observed that some TrpB variants lose binding affinity for PLP resulting in non-functional apoenzyme. We evaluated Trp formation of B5 supplemented with 0, 0.1, 0.25, 0.5, 1, 2, 5, and 100 μM PLP via UV-Vis spectrophotometry. Serine (final conc. 25 mM) + PLP master mixes of the eight concentrations were prepared and dispensed into 96-well UV-transparent plate. Enzyme (final conc. 1 μM) with or without indole master mixes were prepared and 100 µL dispensed into 96-well plate. The plate was immediately transferred into plate reader, shaken for 10 sec to mix and product formation was measured ~20 sec for 120 cycles at 290 nm.

Only the 100 µM condition restored activity, supporting our hypothesis that the purified enzyme was apoprotein and binds PLP poorly, requiring supplementation of PLP to re-form a functional holoenzyme. Thus, we chose to supplement PLP in the subsequent purified protein reactions.

### B5 and G6 small-scale analytical reactions

Reactions were performed in 2 mL glass HPLC vials (Agilent) charged with nucleophile (final conc. 20 mM) dissolved in DMSO (5% w/v), followed by the addition of purified protein (final enzyme conc. either 2 μM or 40 μM), PLP (final conc. 100 μM) and serine (final conc. 20 mM) to achieve a final volume of 200 μL. Reactions were incubated at 30 °C for ~24 hours. Reactions were diluted with 600 μL 2:1 CH_3_CN/1 M aq. HCl, subjected to centrifugation at 5,000*g*, and 400 μL transferred to 2 mL glass HPLC vials (Agilent). Samples were analyzed by HPLC-MS. Azulene samples were further diluted 20X to avoid oversaturation of the UV-detector and analyzed via UHPLC-MS.

## Supplementary Information for

### Supplementary Tables

**Supplementary Table 1.**
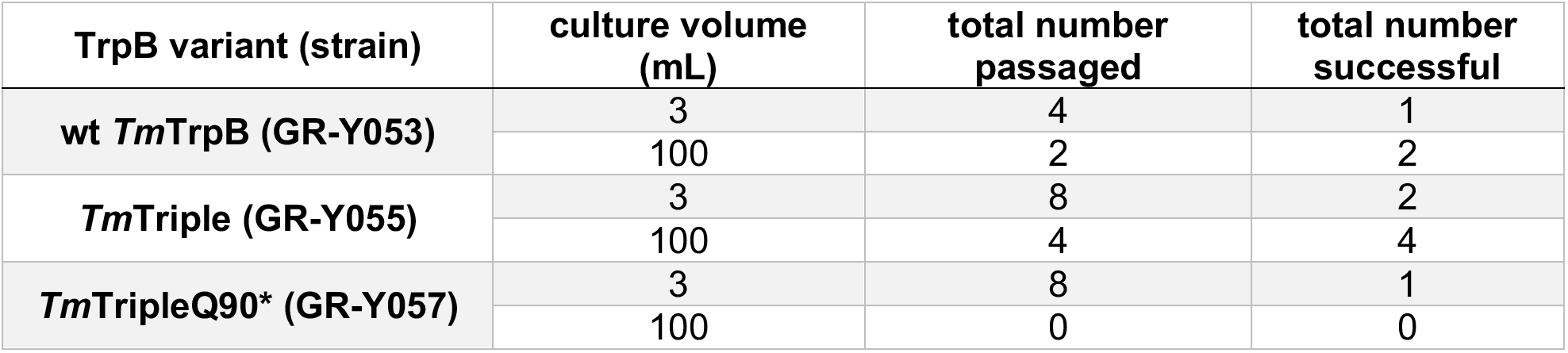
Summary of all cultures passaged for evolution of *Tm*TrpB variants.

| TrpB variant (strain) | culture volume (mL) | total number passaged | total number successful |
| --- | --- | --- | --- |
| **wt *Tm*TrpB (GR-Y053)** | 3 | 4 | 1 |
|  | 100 | 2 | 2 |
| ***Tm*Triple (GR-Y055)** | 3 | 8 | 2 |
|  | 100 | 4 | 4 |
| ***Tm*TripleQ90\* (GR-Y057)** | 3 | 8 | 1 |
|  | 100 | 0 | 0 |

**Supplementary Table 2.** Mutation summary statistics for OrthoRep-evolved TrpB populations.

|  |  | non-synonymous mutations |  |  | synonymous mutations |  |
| --- | --- | --- | --- | --- | --- | --- |
| variant set | total number of sequences | total unique mutations | mean | standard deviation | mean | standard deviation |
| consensus | 10 | 43 | 5.6 | 2.3 | 3.0 | 2.3 |
| 1 | 16 | 85 | 8.7 | 2.1 | 6.3 | 4.4 |
| 2 | 60 | 194 | 9.3 | 2.8 | 6.5 | 3.0 |

**Supplementary Table 3.** Mutations and identification information for all individual *Tm*TrpB sequences. (See Supplementary_Table_3_Extended.xls for mutations and sequences.)

| variant set | variant ID | variant name | evolution population | number of non-synonymous mutations | number of synonymous mutations | starting sequence |
| --- | --- | --- | --- | --- | --- | --- |
| 1 | A | Tm11g | Tm11 | 13 | 3 | TmTrpB |
| 1 | B | Tm12f | Tm12 | 9 | 7 | TmTrpB |
| 1 | C | Tm1Ff | Tm1F | 7 | 1 | TmTrpB |
| 1 | D | Tm4Ge | Tm4G | 6 | 11 | TmTripleQ90* |
| 1 | E | Tm4Gf | Tm4G | 11 | 15 | TmTripleQ90* |
| 1 | F | Tm3Cc | Tm3C | 9 | 3 | TmTriple |
| 1 | G | Tm3Cf | Tm3C | 9 | 3 | TmTriple |
| 1 | H | Tm3Cg | Tm3C | 11 | 4 | TmTriple |
| 1 | R | Tm3Df | Tm3D | 9 | 6 | TmTriple |
| 1 | J | Tm33f | Tm33 | 10 | 5 | TmTriple |
| 1 | K | Tm34e | Tm34 | 9 | 7 | TmTriple |
| 1 | L | Tm34f | Tm34 | 8 | 6 | TmTriple |
| 1 | M | Tm35f | Tm35 | 5 | 3 | TmTriple |
| 1 | N | Tm36b | Tm36 | 7 | 11 | TmTriple |
| 1 | O | Tm36e | Tm36 | 10 | 1 | TmTriple |
| 1 | P | Tm36f | Tm36 | 6 | 14 | TmTriple |
| 2 | B1 | Tm11-B1 | Tm11 | 16 | 11 | wt TmTrpB |
| 2 | B3 | Tm11-B3 | Tm11 | 12 | 7 | wt TmTrpB |
| 2 | B4 | Tm11-B4 | Tm11 | 14 | 10 | wt TmTrpB |
| 2 | B6 | Tm11-B6 | Tm11 | 14 | 9 | wt TmTrpB |
| 2 | B8 | Tm11-B8 | Tm11 | 12 | 9 | wt TmTrpB |
| 2 | D5 | Tm12-D5 | Tm12 | 9 | 10 | wt TmTrpB |
| 2 | D4 | Tm12-D4 | Tm12 | 7 | 5 | wt TmTrpB |
| 2 | D8 | Tm12-D8 | Tm12 | 7 | 6 | wt TmTrpB |
| 2 | D7 | Tm12-D7 | Tm12 | 9 | 8 | wt TmTrpB |
| 2 | D6 | Tm12-D6 | Tm12 | 10 | 5 | wt TmTrpB |
| 2 | D3 | Tm12-D3 | Tm12 | 7 | 9 | wt TmTrpB |
| 2 | C2 | Tm12-C2 | Tm12 | 9 | 9 | wt TmTrpB |
| 2 | D1 | Tm12-D1 | Tm12 | 8 | 9 | wt TmTrpB |
| 2 | A1 | Tm1F-A1 | Tm1F | 9 | 4 | wt TmTrpB |
| 2 | A7 | Tm1F-A7 | Tm1F | 10 | 8 | wt TmTrpB |
| 2 | A6 | Tm1F-A6 | Tm1F | 8 | 4 | wt TmTrpB |
| 2 | A5 | Tm1F-A5 | Tm1F | 8 | 6 | wt TmTrpB |
| 2 | A4 | Tm1F-A4 | Tm1F | 7 | 5 | wt TmTrpB |
| 2 | A3 | Tm1F-A3 | Tm1F | 7 | 7 | wt TmTrpB |
| 2 | A2 | Tm1F-A2 | Tm1F | 8 | 4 | wt TmTrpB |
| 2 | A8 | Tm1F-A8 | Tm1F | 6 | 2 | wt TmTrpB |
| 2 | F3 | Tm4G-F3 | Tm4G | 14 | 8 | TmTripleQ90* |
| 2 | F5 | Tm4G-F5 | Tm4G | 14 | 8 | TmTripleQ90* |
| 2 | F6 | Tm4G-F6 | Tm4G | 11 | 9 | TmTripleQ90* |
| 2 | F7 | Tm4G-F7 | Tm4G | 9 | 6 | TmTripleQ90* |
| 2 | F2 | Tm4G-F2 | Tm4G | 16 | 6 | TmTripleQ90* |
| 2 | F1 | Tm4G-F1 | Tm4G | 11 | 9 | TmTripleQ90* |
| 2 | C8 | Tm3C-C8 | Tm3C | 10 | 5 | TmTriple |
| 2 | C5 | Tm3C-C5 | Tm3C | 15 | 6 | TmTriple |
| 2 | D2 | Tm3C-D2 | Tm3C | 9 | 4 | TmTriple |
| 2 | C1 | Tm3C-C1 | Tm3C | 6 | 4 | TmTriple |
| 2 | C7 | Tm3C-C7 | Tm3C | 12 | 3 | TmTriple |
| 2 | C3 | Tm3C-C3 | Tm3C | 7 | 5 | TmTriple |
| 2 | E8 | Tm3D-E8 | Tm3D | 8 | 11 | TmTriple |
| 2 | E3 | Tm3D-E3 | Tm3D | 13 | 4 | TmTriple |
| 2 | E1 | Tm3D-E1 | Tm3D | 10 | 3 | TmTriple |
| 2 | E2 | Tm3D-E2 | Tm3D | 12 | 8 | TmTriple |
| 2 | G1 | Tm33-G1 | Tm33 | 5 | 5 | TmTriple |
| 2 | G2 | Tm33-G2 | Tm33 | 9 | 6 | TmTriple |
| 2 | G3 | Tm33-G3 | Tm33 | 6 | 5 | TmTriple |
| 2 | G4 | Tm33-G4 | Tm33 | 10 | 8 | TmTriple |
| 2 | G7 | Tm33-G7 | Tm33 | 7 | 2 | TmTriple |
| 2 | G5 | Tm33-G5 | Tm33 | 8 | 4 | TmTriple |
| 2 | G6 | Tm33-G6 | Tm33 | 6 | 5 | TmTriple |
| 2 | H3 | Tm34-H3 | Tm34 | 8 | 5 | TmTriple |
| 2 | H6 | Tm34-H6 | Tm34 | 10 | 15 | TmTriple |
| 2 | H4 | Tm34-H4 | Tm34 | 12 | 10 | TmTriple |
| 2 | H7 | Tm34-H7 | Tm34 | 10 | 10 | TmTriple |
| 2 | H2 | Tm34-H2 | Tm34 | 13 | 4 | TmTriple |
| 2 | H1 | Tm34-H1 | Tm34 | 9 | 15 | TmTriple |
| 2 | H5 | Tm34-H5 | Tm34 | 9 | 10 | TmTriple |
| 2 | C6 | Tm35-C6 | Tm35 | 6 | 2 | TmTriple |
| 2 | B2 | Tm35-B2 | Tm35 | 8 | 4 | TmTriple |
| 2 | B5 | Tm35-B5 | Tm35 | 5 | 2 | TmTriple |
| 2 | B7 | Tm35-B7 | Tm35 | 6 | 3 | TmTriple |
| 2 | C4 | Tm35-C4 | Tm35 | 7 | 3 | TmTriple |
| 2 | E5 | Tm36-E5 | Tm36 | 6 | 8 | TmTriple |
| 2 | E7 | Tm36-E7 | Tm36 | 8 | 7 | TmTriple |
| 2 | E4 | Tm36-E4 | Tm36 | 6 | 6 | TmTriple |
| 2 | F8 | Tm36-F8 | Tm36 | 7 | 5 | TmTriple |

**Supplementary Table 4.** Kinetic parameters of selected TmTrpB variants at 30 °C.

| **variant** | **$k_{\text{cat}}$ [95% credible region] ($\text{s}^{-1}$)** | **$K_{\text{M}}$ [95% credible region] ($\mu\text{M}$)** | **$k_{\text{cat}}/K_{\text{M}}$ [95% credible region] ($\text{mM}^{-1} \text{s}^{-1}$)** |
| --- | --- | --- | --- |
| ***Tm*Triple** | 0.2 [0.16, 0.31] | 41.23 [14.32, 192.66] | 4.89 [1.54, 12.12] |
| **C** | 0.53 [0.49, 0.58] | 3.89 [1.85, 7.99] | 137.22 [70.29, 276.04] |
| **D** | 0.77 [0.72, 0.83] | 5.79 [3.82, 8.8] | 133.38 [91.24, 193.71] |
| **K** | 0.62 [0.59, 0.66] | 5.58 [3.99, 7.91] | 111.89 [81.52, 152.25] |

### Supplementary Figures

**Supplementary Fig. 1.**
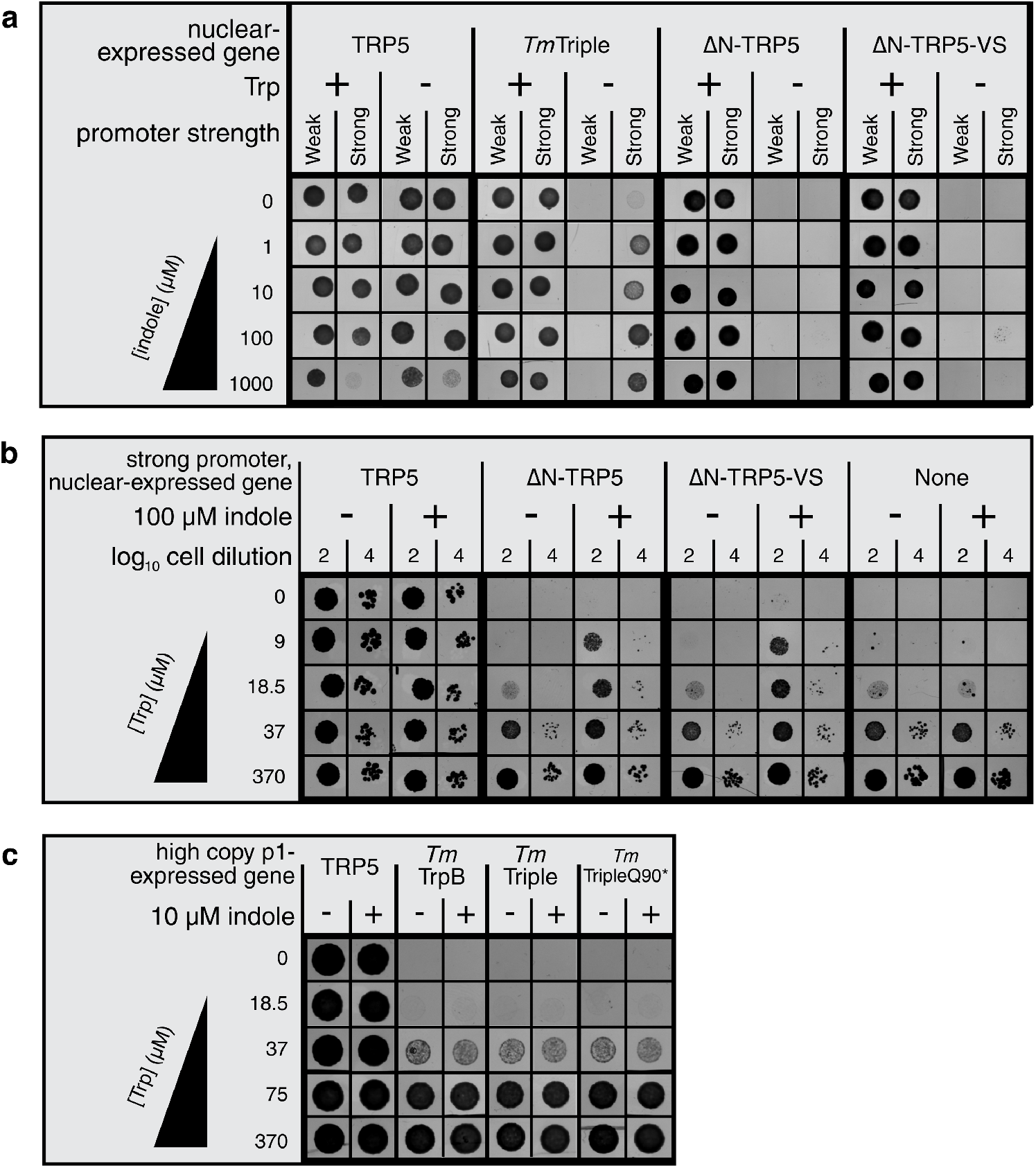
Evaluation of indole-dependent TRP5 complementation of TrpB variants. **a-c**, Spot plating assays for Δ*trp5* yeast strains expressing TrpB variants from a nuclear plasmid under two different promoter strengths (**a**), from a nuclear plasmid under a strong promoter (**b**), or from p1 at a high copy number (wt TP-DNAP1 expressed *in trans*) (**c**) grown on indicated growth medium.

**Supplementary Fig. 2.**
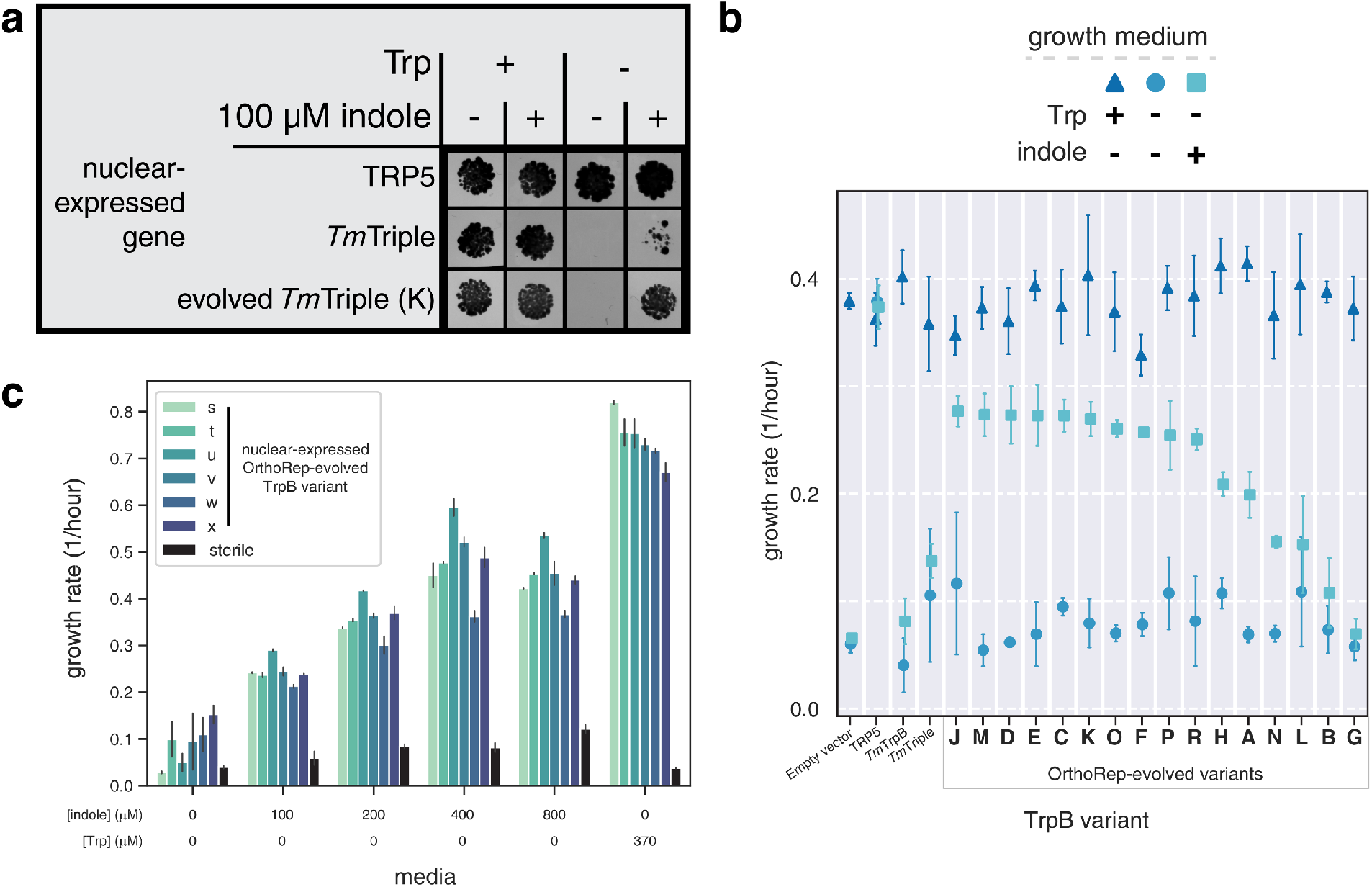
*In vivo* Trp production by evolved TrpBs. **a**, Spot plating assay for TRP5-deleted yeast with a nuclear plasmid expressing TRP5, *Tm*Triple, or an individual OrthoRep-evolved *Tm*TrpB variant driven by a promoter (pRNR2) that approximates expression of *Tm*TrpB variants from OrthoRep’s p1 plasmid, grown on indicated media. **b**, Evaluation of TRP5 complementation by evolved variants through a growth rate assay. Maximum growth rates over a 24-hour period for *Δtrp5* yeast strains transformed with a nuclear plasmid expressing the indicated *Tm*TrpB variant, grown in medium with or without 100 μM indole, and with or without Trp, as indicated. Points and error bars represent mean ± s.d. of four technical replicates, respectively. **c**, Optimization of indole concentration. Growth rates during exponential growth for Δ*trp5* yeast with a nuclear plasmid expressing randomly chosen OrthoRep-evolved TrpB variants (exact sequence not determined), supplemented with one of four indole concentrations, Trp, or neither of these. Bars and error bars represent mean ± s.d. of four technical replicates, respectively.

**Supplementary Fig. 3.**
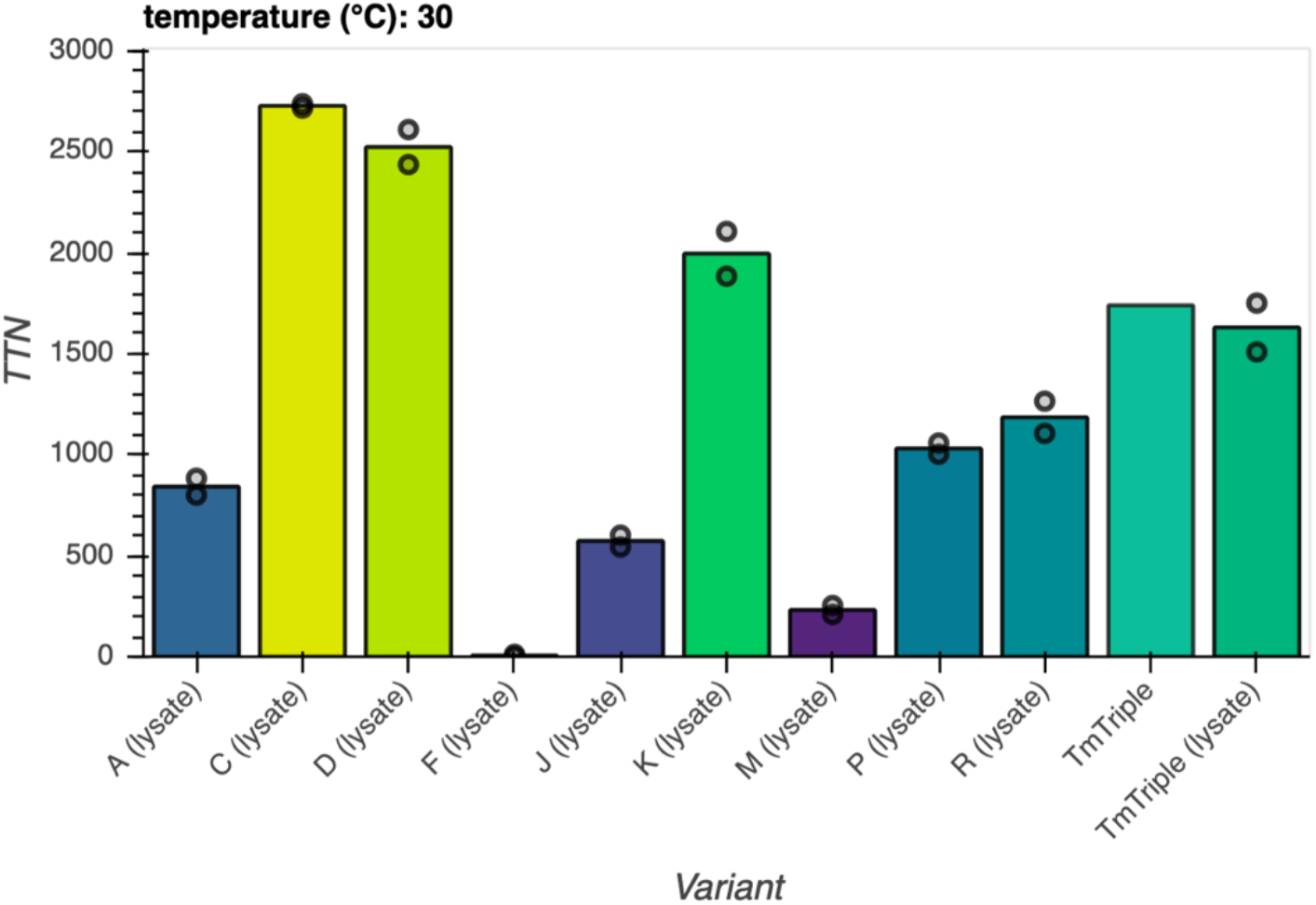
*In vitro* Trp production by evolved TrpBs with heat treated lysate. Trp production at 30 °C by indicated *Tm*TrpB variants. Reactions with *Tm*Triple were performed with both heat treated lysate and with purified protein, while all other reactions were performed only with heat treated lysate. TTN, total turnover number. Maximum TTN is 5,000. Points represent TTN for individual replicates, bars represent mean TTN for reactions with replicates, or TTN for a single replicate otherwise.

**Supplementary Fig. 4.**
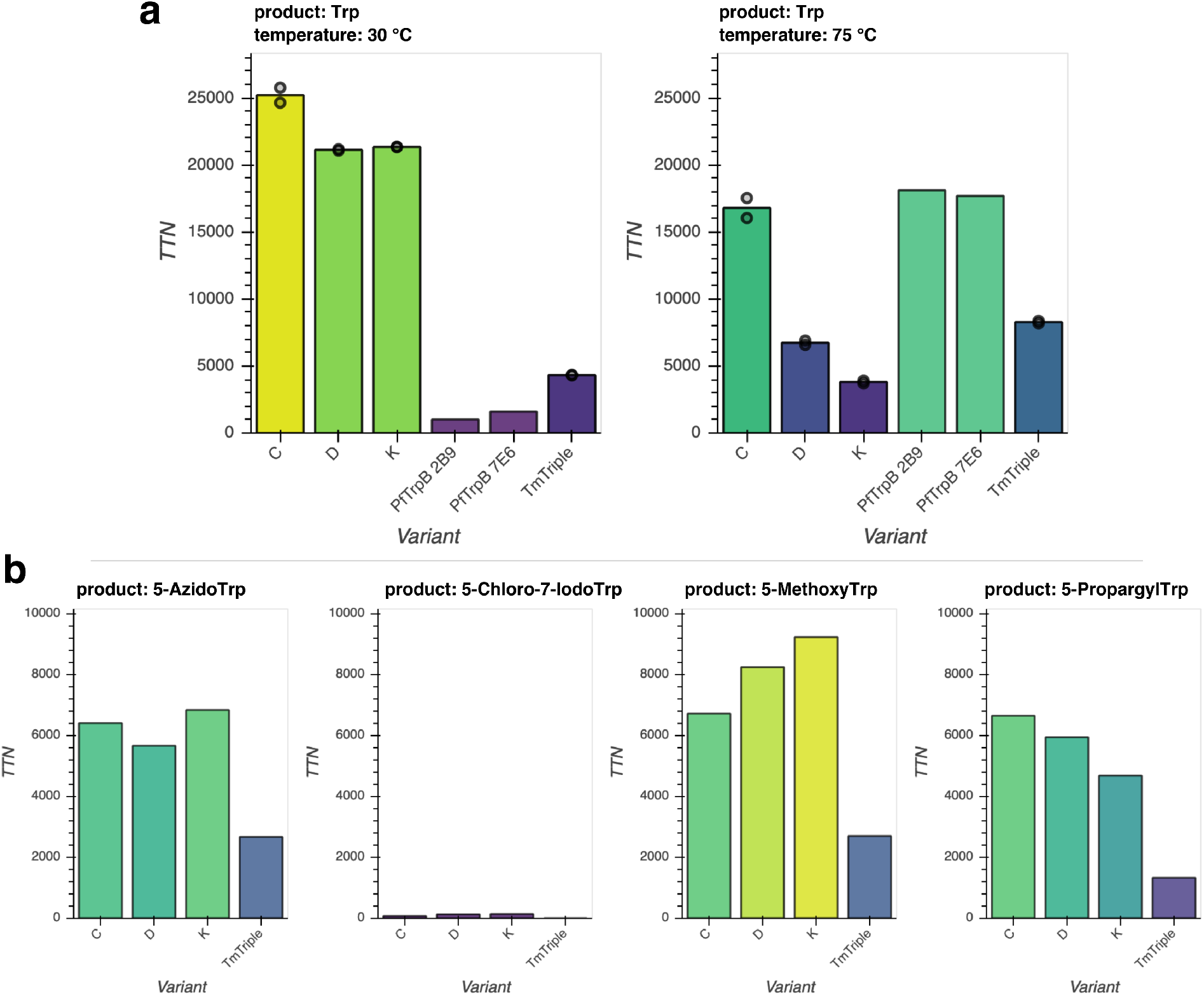
*In vitro* Trp and Trp analog production with purified enzyme. **a**,**b**, Production of (**a**) Trp at 30 °C or 75 °C with 40,000 maximum TTN or (**b**) indicated Trp analogs at 30 °C by column purified *Tm*TrpB variants. TTN, total turnover number with 10,000 as maximum TTN. Points represent TTN for individual replicates, bars represent mean TTN for reactions with replicates, or TTN for a single replicate otherwise.

**Supplementary Fig. 5.**
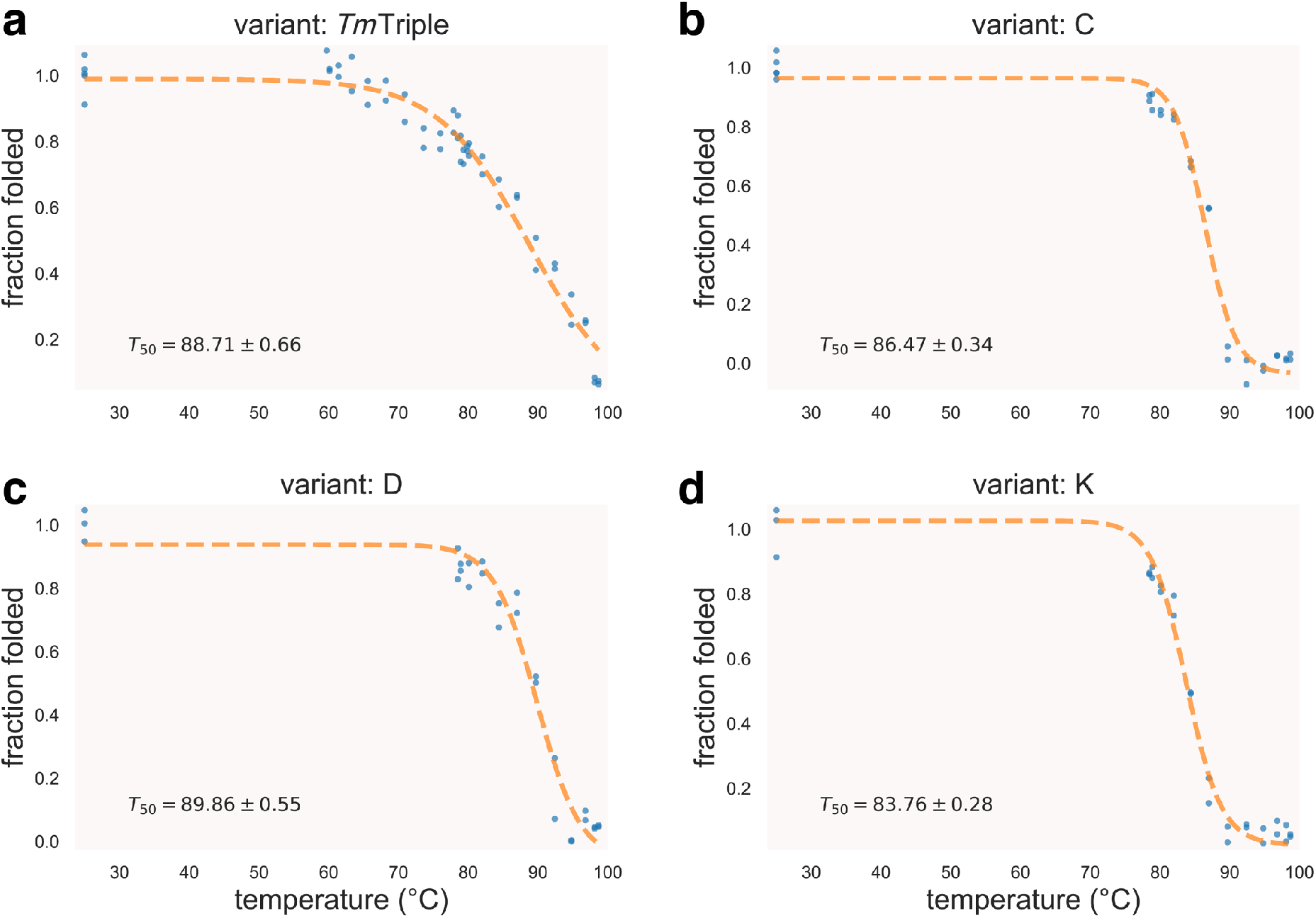
Thermal shift assay on various *Tm*TrpBs. Proportion of *Tm*TrpB variants TmTriple (**a**), C (**b**), D (**c**), or K (**d**) that remain folded after incubation at indicated temperature for 1 hour, as measured by the fraction of Trp production relative to incubation at 25 °C. *T*_50_, temperature at which 50% of enzyme is irreversibly inactivated, as estimated by best fit logistic model (dotted line). Each temperature tested in duplicate.

**Supplementary Fig. 6.**
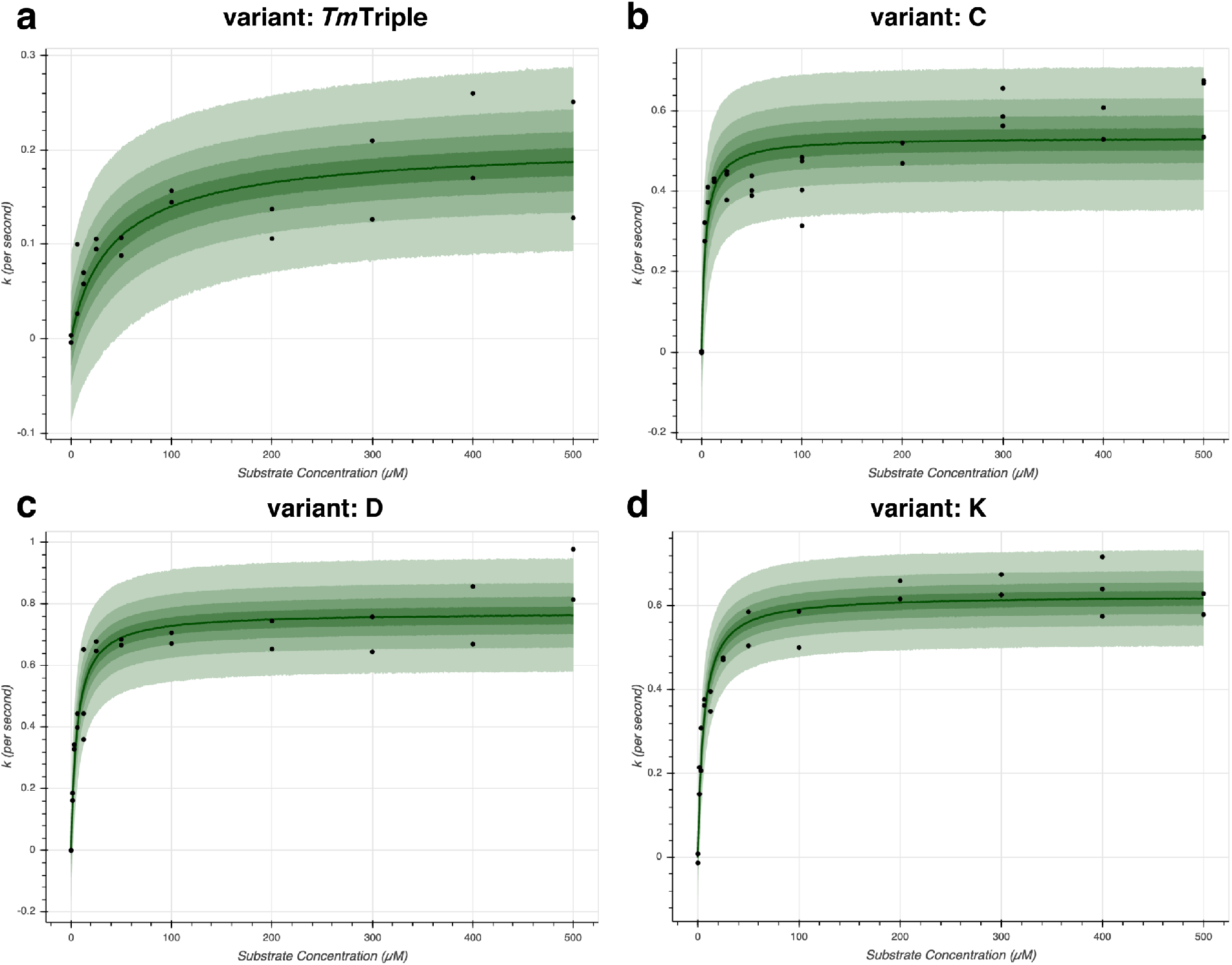
Michaelis-Menten plots for rate of Trp production at saturating serine for evolved *Tm*TrpB variants. Initial rate of Trp formation (*k*, per second) with TrpB variants TmTriple (**a**), C (**b**), D (**c**), or K (**d**) at saturating serine concentration (40 mM) vs. indole concentration. Points, median estimates for initial rate based on absorbance change over time (see **Methods**). The median estimated Michaelis-Menten curve is shown as a dark green line, with the 25, 50, 75, and 95% credible regions displayed from dark to light green, respectively. All measurements performed in at least duplicate.

**Supplementary Fig. 7.**
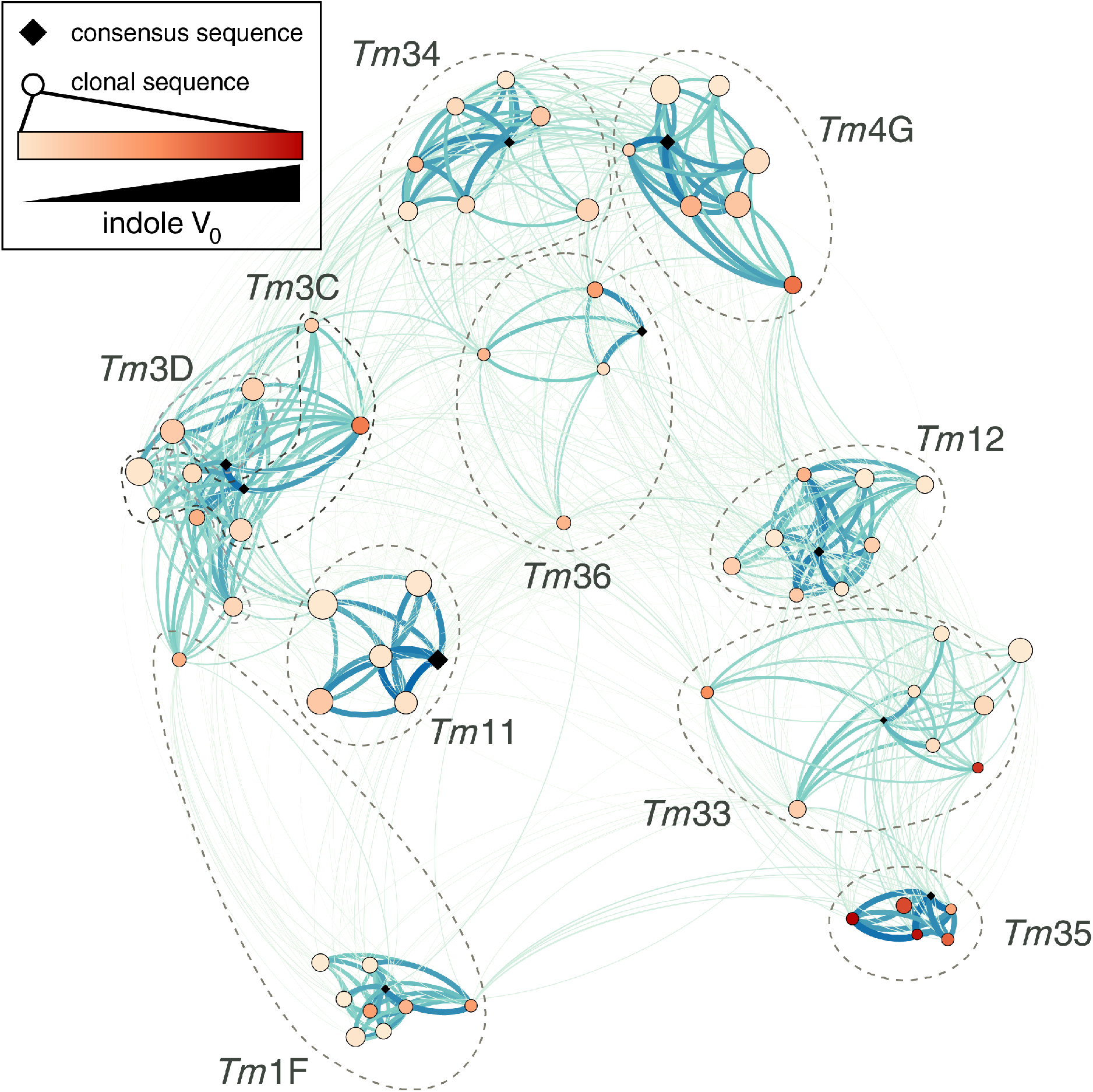
Relatedness of TrpB panel sequences generated by OrthoRep evolution. Force directed graph where each node represents an individual sequence (all variants from set 2, and variants **C**, **D**, and **K**) or consensus sequence for one of the ten evolved populations. Edge weights are proportional to the number of shared mutations between two nodes. Higher edge weight yields a stronger attractive force between two nodes, and is visualized as a darker color and a thicker line. Nodes for individual sequences are colored according to initial rate of Trp formation, similar to Fig. 3b. Dotted lines are drawn around consensus sequences and individual sequences that are derived from the same evolved culture, if nodes are sufficiently clustered to allow it.

**Supplementary Fig. 8.**
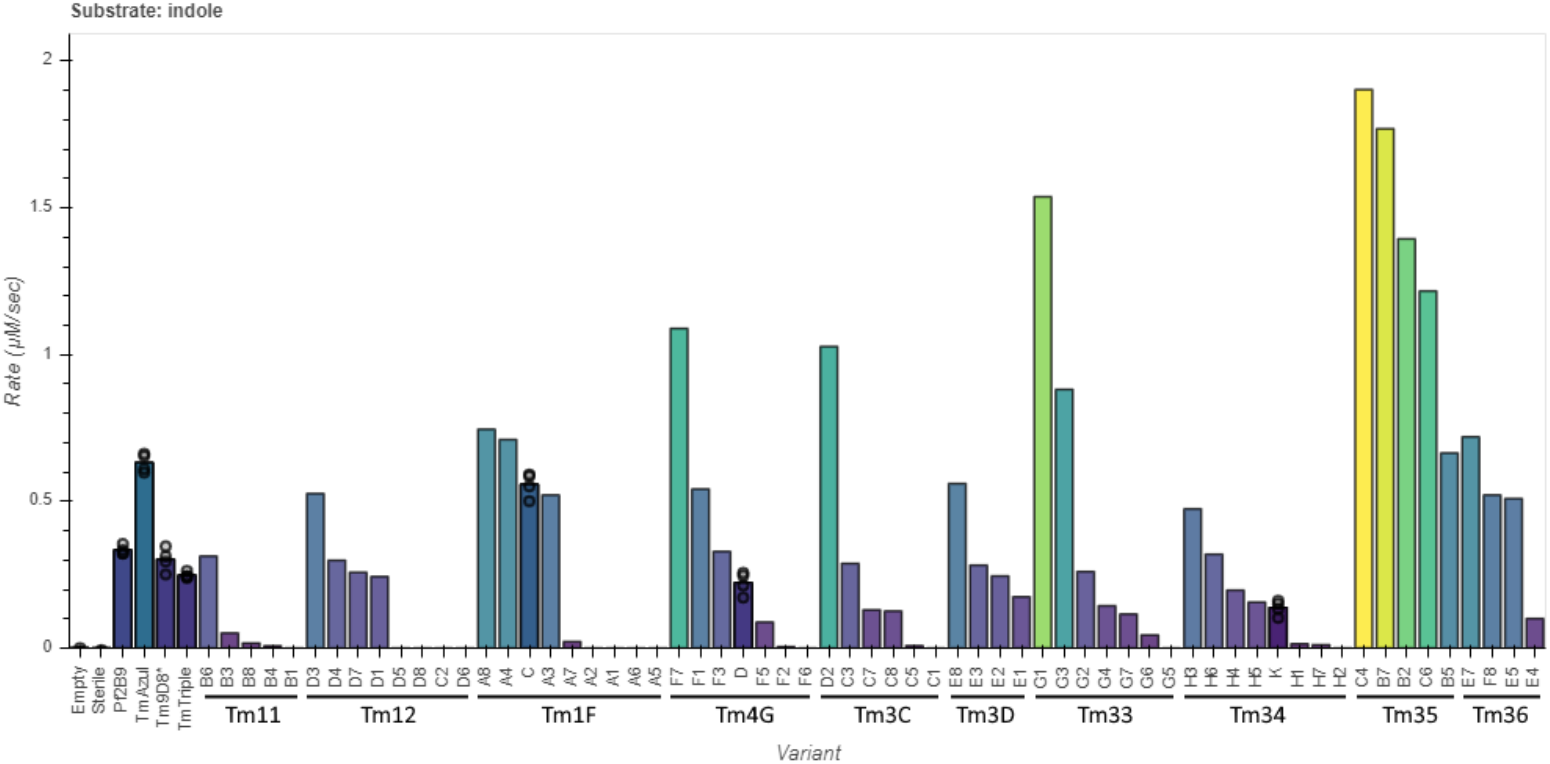
TrpB panel indole activity by initial rate of Trp formation. Initial rate of Trp formation at saturating L-serine by UV-vis spectrophotometry. Points represent rate for individual replicates, bars represent mean rate for reactions with multiple replicates, or rate for a single replicate otherwise. OrthoRep-evolved variants are ordered first by the population from which they were derived, then by indole activity. Empty, expression vector without TrpB variant. Sterile, reaction master mix without heat-treated lysate added. Empty, *Pf*2B9, *Tm*Azul, *Tm*9D8*, *Tm*Triple, **C**, **D**, and **K** all performed in quadruplicate; sterile performed in duplicate; all other reactions performed in a single replicate.

**Supplementary Fig. 9.**
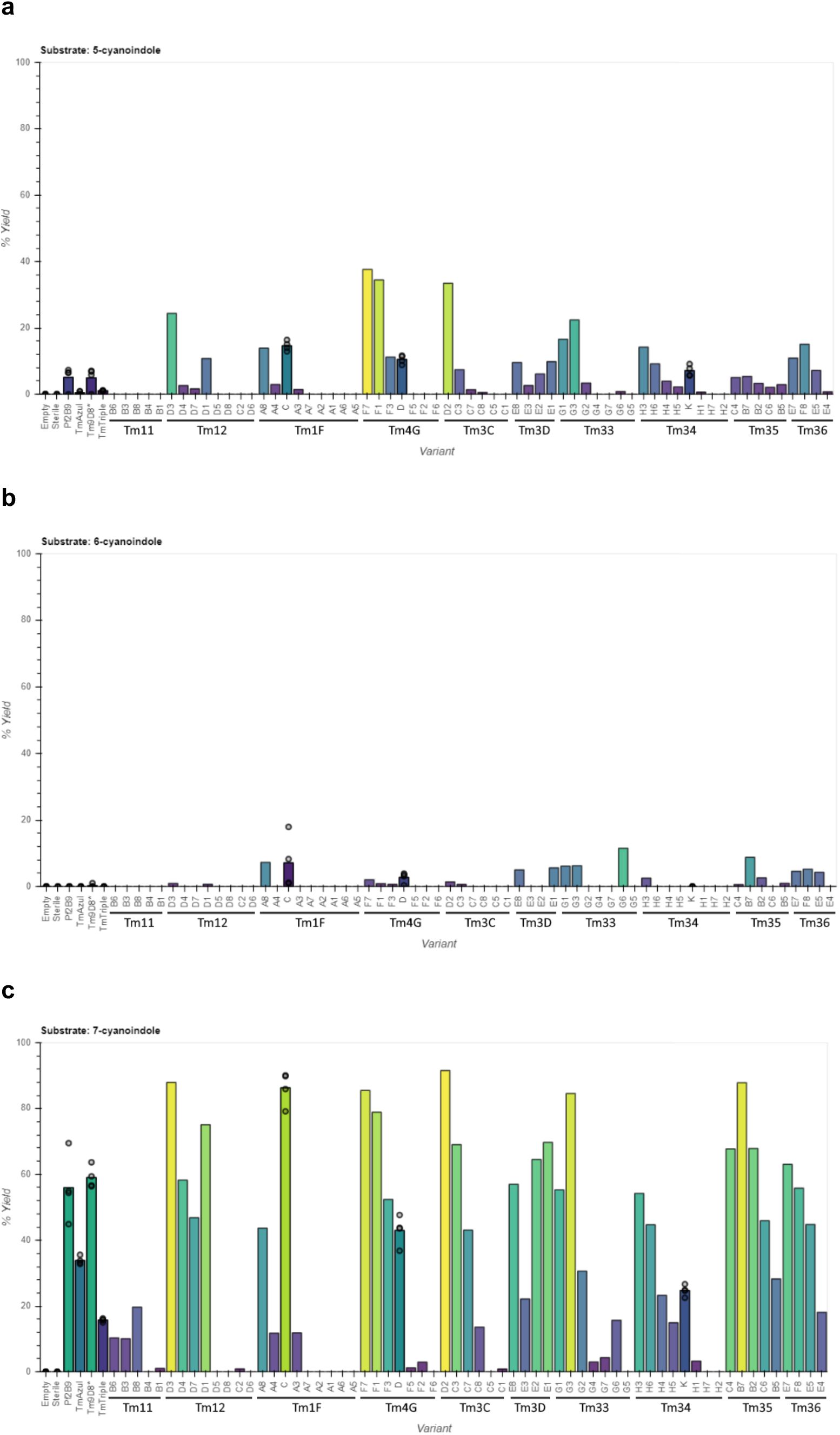
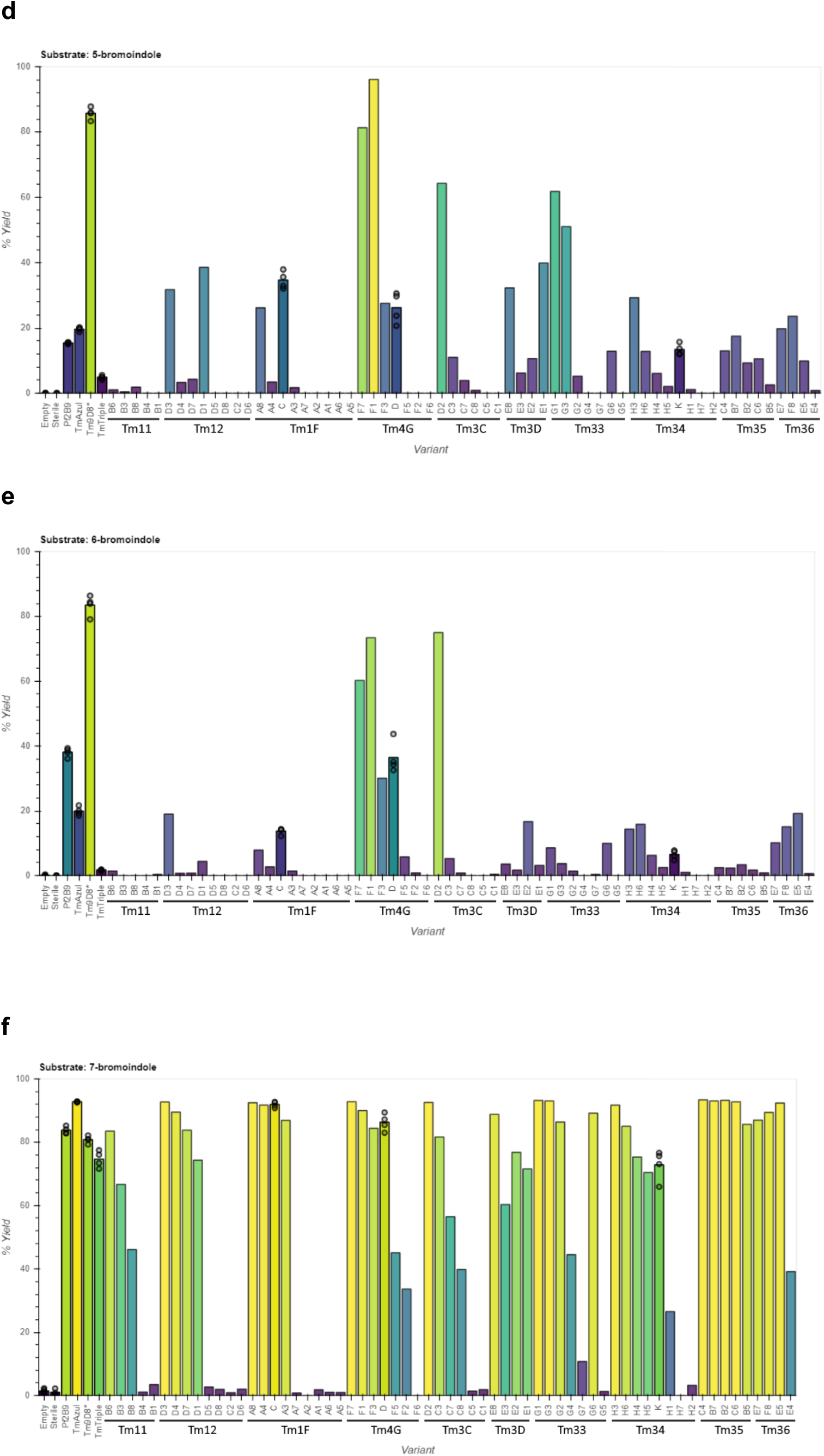
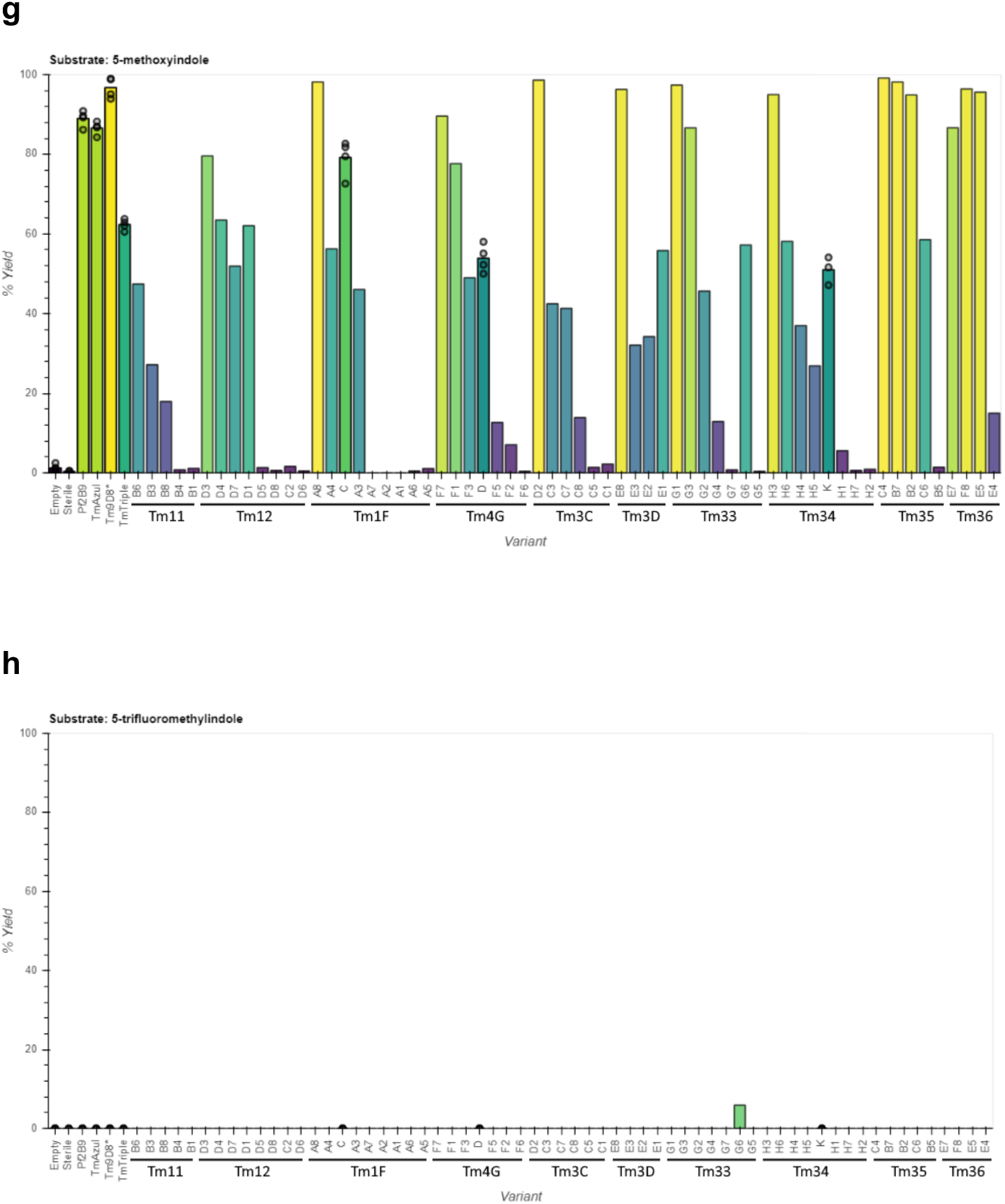
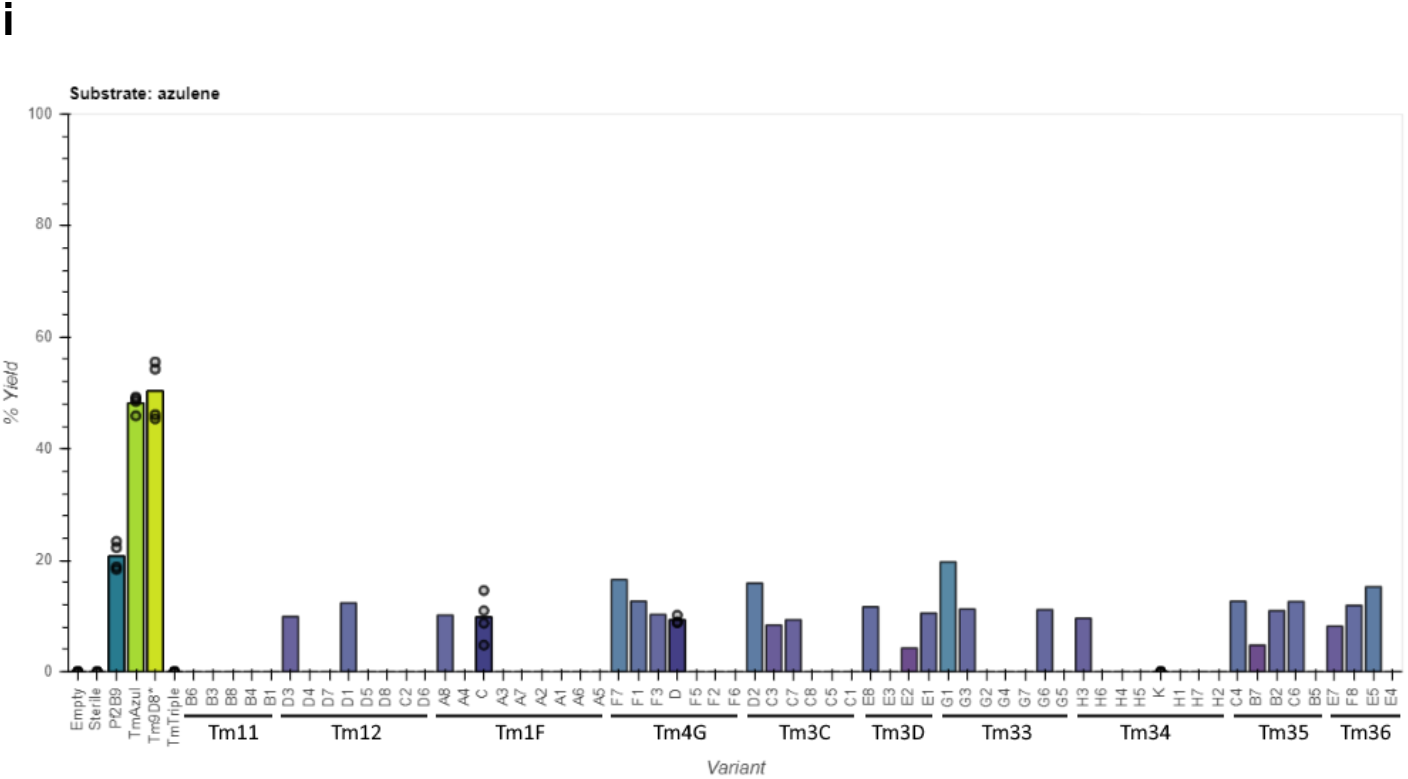
TrpB panel activity with indole analogs by HPLC yield. **a**-**i**, HPLC yield of (**a**) 5-cyanoTrp, (**b**) 6-cyanoTrp, (**c**) 7-cyanoTrp, (**d**) 5-bromoTrp, (**e**) 6-bromoTrp, (**f**) 7-bromoTrp, (**g**) 5-methoxyTrp, (**h**) 5-trifluoromethylTrp, and (**i**) β-(1-azulenyl)-L-alanine for indicated variants supplied with L-serine and each indole substrate. Points represent % yield for individual replicates, bars represent mean % yield for reactions with replicates, or % yield for a single replicate otherwise. Replicates and variant order are as in Fig. S8, and populations from which OrthoRep-evolved variants were derived are annotated.

**Supplementary Fig. 10.**
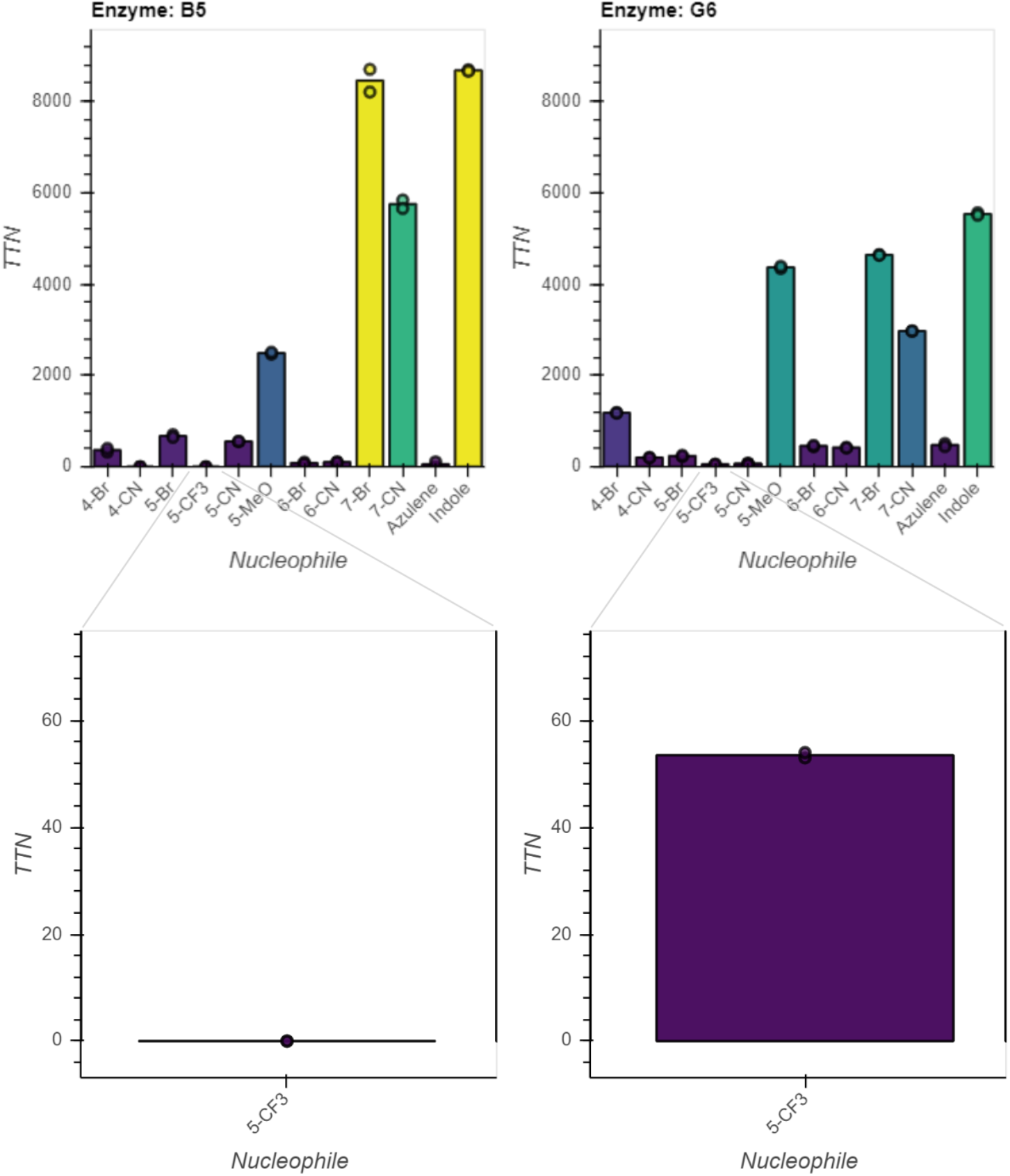
Substrate activity profiles for large scale purification of variants B5 and G6. Total turnover number (TTN) for *Tm*TrpB variants B5 and G6 purified at large scale (see **Methods**) and supplied with L-serine and the indicated indole analog, azulene, or indole (nucleophile), with a maximum TTN of 10,000. All reactions were performed in duplicate. Points represent TTN for individual replicates, bars represent mean for two replicates. Insets, TTN for 5-trifluoromethylindole with y-axis scale adjusted for clarity.

**Supplementary Fig. 11.**
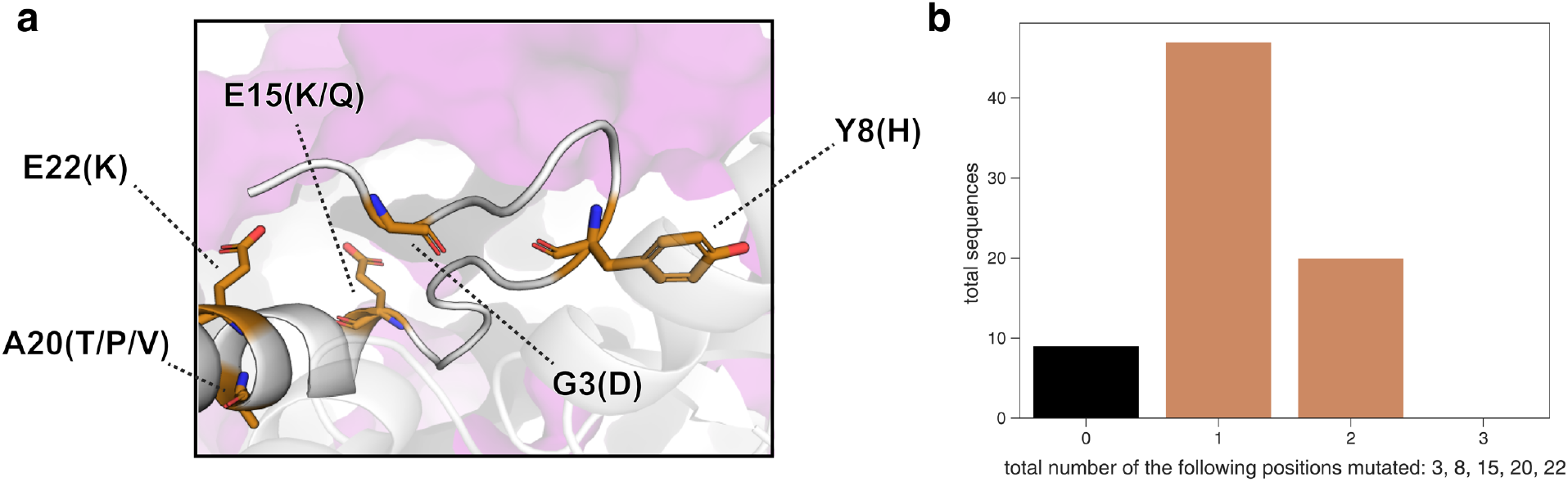
Commonly observed mutations at the α-subunit interaction interface. **a**, Homology-predicted *Tm*TrpB structure (based on engineered stand-alone *Pf*TrpB, PDB 6AM8), with commonly mutated *Tm*TrpB residues located near the TrpA interaction interface highlighted. Solvent-exposed regions of TrpA (purple) (PDB 1WDW) are shown as a surface. Mutations are indicated by the wt residue and position, followed by any residues to which this wt residue is mutated in OrthoRep-evolved TrpB sequences. **b**, Total number of sequences in both variant sets 1 and 2 that contain the indicated number of mutations to any of the residues highlighted in panel **a**.

**Supplementary Fig. 12.**
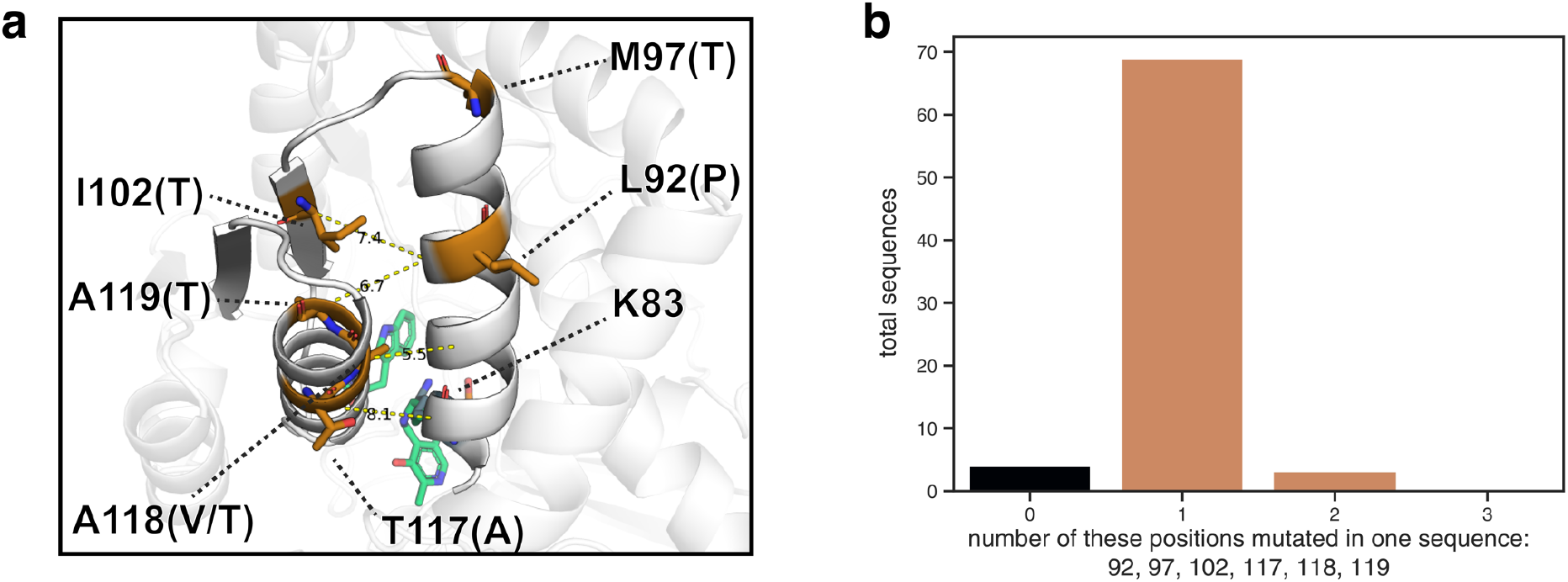
Commonly observed mutations to residues near a catalytic α-helix. **a**, Homology-predicted *Tm*TrpB structure (aligned to engineered stand-alone *Pf*TrpB, PDB 6AM8) with wt residues on or near the α-helix housing K83, which are commonly mutated in OrthoRep evolved populations (orange). PLP (green) and Trp (green) are shown as sticks, and the catalytic lysine K83 (teal) is shown as spheres. Mutations are indicated by the wt residue and position, followed by any residues to which this wt residue is mutated in OrthoRep-evolved TrpB sequences. Dotted lines connect the α-carbon of residues not located on the K83 α-helix with the α-carbon of the nearest residue on the K83 α-helix, with the distance noted in Ångstroms. **b**, Total number of sequences in both variant sets 1 and 2 that contain the indicated number of mutations to any of the residues highlighted in panel **a**.

**Supplementary Fig. 13.**
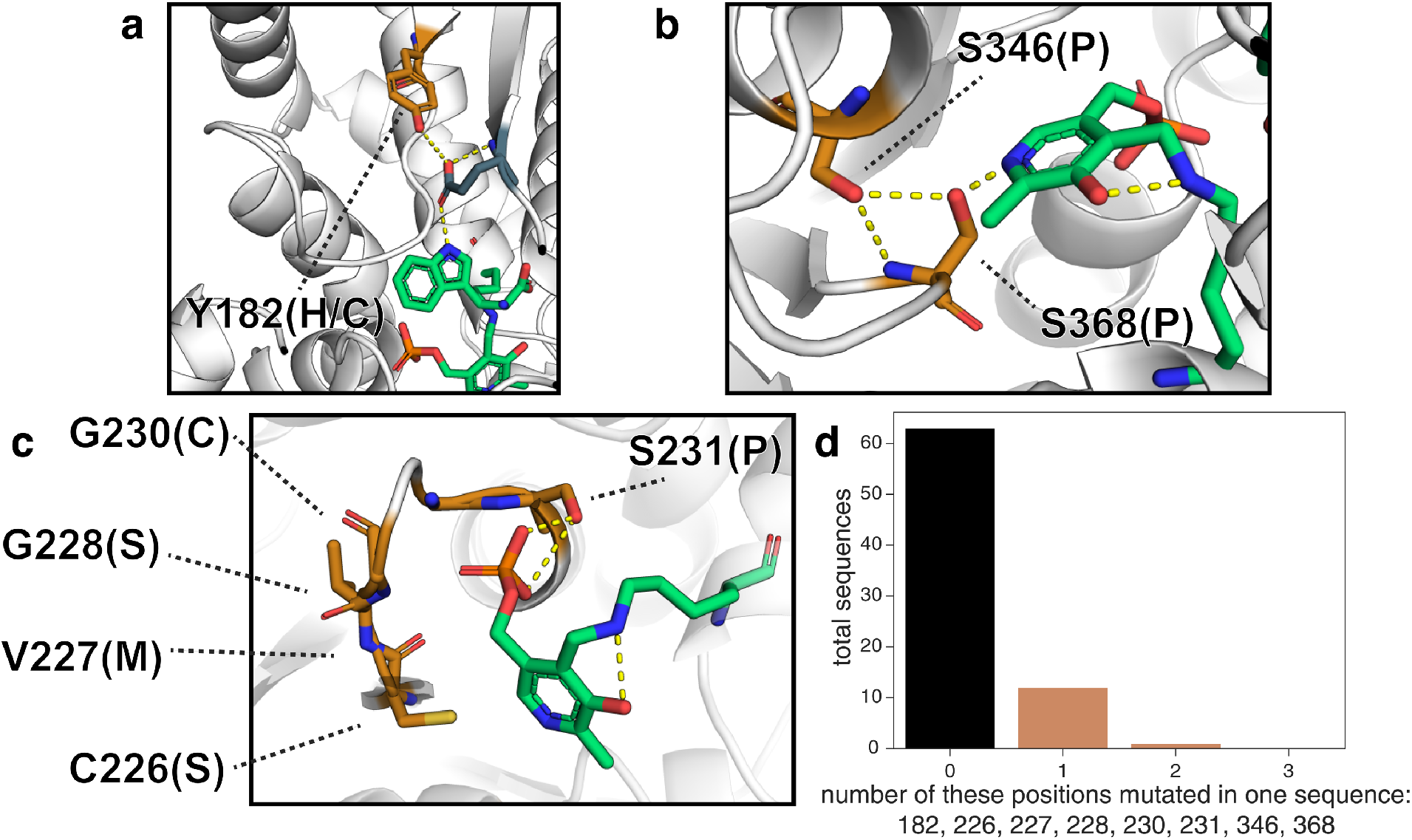
First- and second-shell active site mutations. **a**-**c**, Homology model of *Tm*TrpB (aligned to *Pf*TrpB, PDB: 6AM8) highlighting residues mutated in OrthoRep-evolved variants (orange) that may influence (**a**) indole charge, (**b**) PLP six-member ring binding, and (**c**) PLP-phosphate binding. Mutations are indicated by the wt residue and position, followed by any residues to which this wt residue is mutated in OrthoRep-evolved TrpB sequences. **d**, Total number of sequences in both variant sets 1 and 2 that contain the indicated number of mutations to any of the residues highlighted in panels **a**, **b**, or **c**.

**Supplementary Fig. 14.**
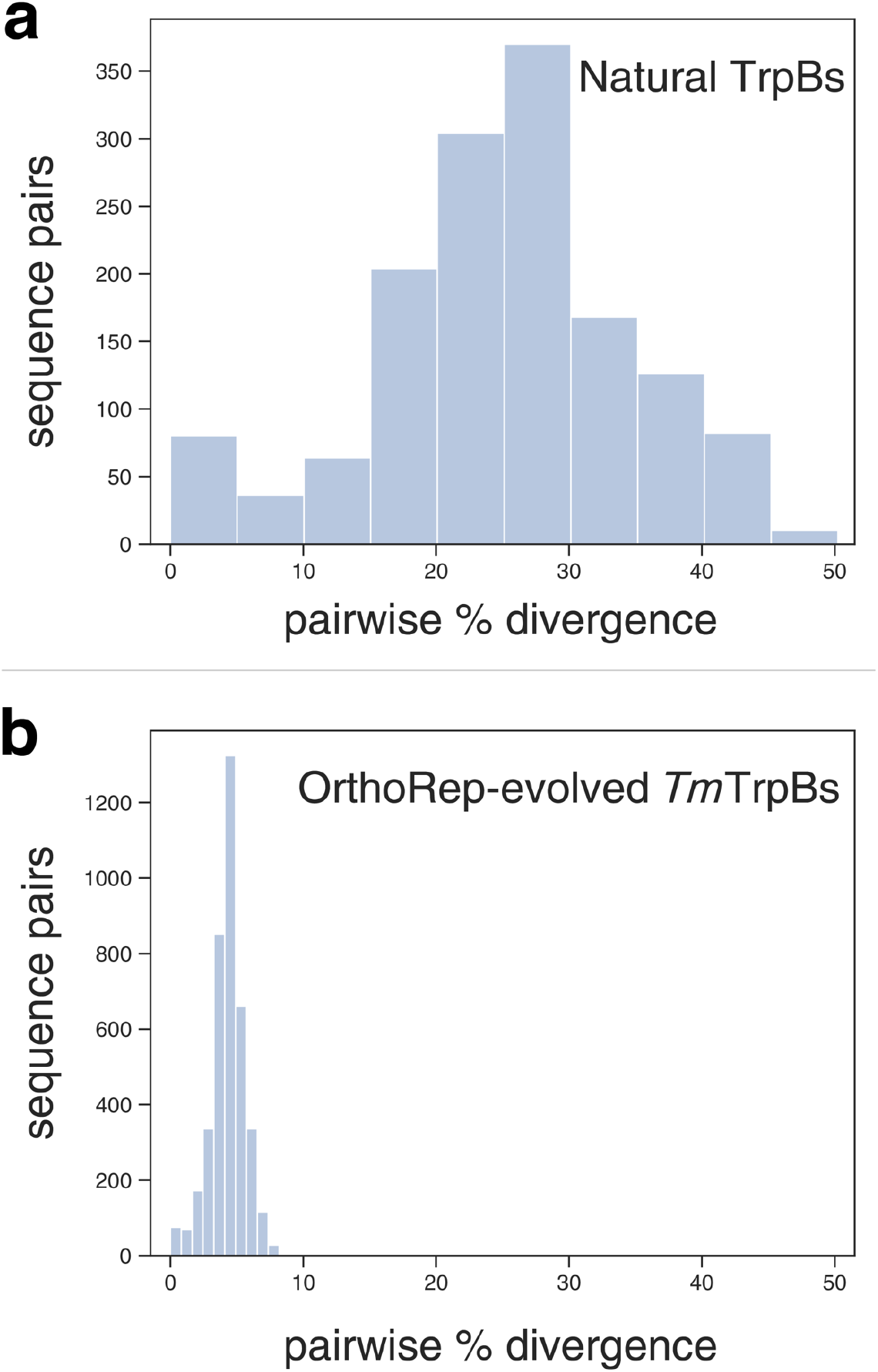
Sequence divergence for natural and OrthoRep-evolved TrpBs. **a-b**, Distributions of pairwise % amino acid sequence divergence for a diverse group of 38 naturally occurring mesophilic TrpB variants (**a**) and OrthoRep-evolved variant sets 1 and 2 (**b**).

